# The alternative splicing of *ZmHsf23* regulates heat stress tolerance in maize

**DOI:** 10.1101/2024.04.28.591503

**Authors:** Jing Wang, Nan-Nan Song, Qian-Qian Qin, An-Qi Su, Wei-Na Si, Bei-Jiu Cheng, Hai-Yang Jiang

## Abstract

- Heat stress is one of the major threats to maize (*Zea mays*) production globally. Heat shock transcription factors (HSFs) play vital roles in plant heat stress responses. However, the molecular and genetic basis of HSFs in maize thermotolerance remain largely unknown.
- In this study, we reveal that the alternative splicing of *Hsf23* in maize modulates heat stress tolerance. *Hsf23* produces two functional transcripts, the full-length transcript *Hsf23b* and the heat-inducible transcript *Hsf23a*. The two *Hsf23* transcripts differ by the presence of a cryptic mini exon in *Hsf23a*, which is spliced out in *Hsf23b*. Both *Hsf23a* and *Hsf23b* were intensely expressed in response to heat stress.
- The overexpression of *Hsf23b*, not *Hsf23a*, enhanced heat stress tolerance, while loss-of-function mutations of *Hsf23a* and *Hsf23b* exhibited remarkably increased sensitivity to heat stress. Transcriptome analysis revealed that Hsf23b activates broader heat-responsive genes than Hsf23a, and Hsf23a and Hsf23b modulate heat stress response through different downstream targets. Furthermore, Hsf23a physically interacted with Hsf23b and promotes Hsf23b-regulated expression of *sHSP* genes.
- Together, our finding provides new insights into the roles of *ZmHsf23* in the heat tolerance in maize, and presents an important candidate for the genetic improvement of heat-tolerant maize varieties.

## Introduction

Heat stress is a major abiotic stress that severely affects plant growth, development, and crop productivity (Lesk *et al*., 2016; Lohani *et al*., 2020). Maize (*Zea mays*) is an important food crop worldwide, and its growth and development are much susceptible to heat stress (Zhao *et al*., 2017). With the increases in annual global temperature, heat stresses have become a major constraint impacting yields of maize (Zhao *et al*., 2017; Slafer & Savin, 2018). Therefore, it is necessary to study the complex processes and regulation mechanisms of heat stress response (HSR) in maize that may help to identified new heat stress responsive genes and improved maize thermotolerance. Moreover, the understanding of plant HSR systems offer clues and opportunity to develop heat-tolerant maize varieties.

Heat stresses are perceived by various types of plant thermosensors (Lin *et al*., 2020), and the heat stress signals were transmitted to activate downstream stress-transducing transcription factors (TFs) and HS-regulated genes, such as heat shock transcription factors (HSFs) and Heat shock proteins (HSPs) (Fragkostefanakis *et al*., 2015; Li *et al*., 2018). HSFs are considered to play central regulatory roles in the HSR and in the acquisition of thermotolerance in plants (Li *et al*., 2020). They mediate the activation of *HSP* genes and other heat-responsive genes by interacting with the conserved heat stress elements (HSEs) in their promoters in response to heat stress (Guo *et al*., 2016; Ohama *et al*., 2016). Plant HSFs are classified into three classes: HSFA, HSFB and HSFC. HsfAs have short activator motifs (AHA motifs) located in their C-terminal domains, which give HsfAs the ability to function as transcription activators (Guo *et al*., 2016). Among the plant HSFs, HsfA1s have been reported to function as ‘master regulators’ that are essential for the activation of transcriptional networks of HSR (Liu *et al*., 2011). HsfA1s directly regulate the expression levels of many HS-responsive TFs such as DREB2A, HsfA2, HsfA7a, HsfBs and MBF1c (Ohama *et al*., 2016). Loss of function of HsfA1 genes resulted in reduced HS-responsive genes, showing HS-sensitive phenotypes both in tomato and *Arabidopsis* (Chan-Schaminet *et al*., 2009; Andrási *et al*., 2021). HsfA2 has been identified as the dominant HSF in HSR and function downstream of HSFA1s. In addition, HSFA2 can interacts directly with HSFA1s to synergistically activate the expression of HS-regulated genes in response to heat stress (Chan-Schaminet *et al*. 2009; Liu & Charng, 2013). Ectopic expression of lily *HSFA2* and maize *HSFA2* in *Arabidopsis* increased heat stress tolerance of seedling (Xin *et al*., 2010; Gu *et al*., 2019). Besides the HSFA1 and HSFA2, other HSFA members are also involved in the response to heat stress. For instance, lily *HsfA3A* and *HsfA3B* enhanced plant acquired thermotolerance when overexpressed in *Arabidopsis* (Wu *et al*., 2018). Similarly, the overexpression of *TaHsfA5* and *TaHSFA6f* improved the thermotolerance of transgenic plants (Xue *et al*., 2015; Samtani *et al*., 2023).

HSFs are mainly regulated at four levels (Andrási *et al*., 2021). Recent studies suggested that the post-transcriptional regulation of plant HSFs, especially alternative splicing (AS), are essential for plant HSRs (Ling *et al*., 2021). AS produces multiple transcripts from a single pre-mRNA by using different splice sites, thereby enhancing the coding capacity of the genome and increasing the complexity of proteome in response to environmental stresses (Laloum *et al*., 2018; Ling *et al*., 2021). Notably, heat stress markedly enhanced the frequency of AS events in plants, and this could provide a specific means for regulating gene expression in response to heat stress (Jiang *et al*., 2017; Li *et al*., 2021). HS-induced AS of *HSFA2* is observed in *Arabidopsis* and tomato. In *Arabidopsis*, except for full spliced AtHsfA2-I, AtHsfA2-II and AtHsfA2-III were generated due to the different intron retention in the DBD (Liu *et al*., 2013). The HsfA2-II isoform was increased accumulated in wild tomato species, resulting in the improved thermotolerance compared with domesticated varieties (Hu *et al*., 2020). AS induced by HS is also observed for *OsHSFA2d* in rice. OsHSFA2dII is the dominant splice variant of *OsHSFA2d* under normal conditions, however, when exposed to HS conditions, *OsHSFA2d* spliced to generate the active forms OsHSFA2dI, which functions as a transcriptional activator of *HSPs* and *OsBiP1* in HSR (Cheng *et al*., 2015). In addition, *HSF04* and *HSF17* in maize, *HSFA3* in lily, as well as *HSFA6e* in wheat, are also subjected to alternative splicing in response to heat stress (Wu *et al*., 2019; Zhang *et al*., 2020; Wen *et al*., 2023).

In this study, we aimed to explore the functions of Hsf23a and Hsf23b, two splice variants of *Hsf23*, in heat stress response in maize. We revealed that both *Hsf23a* and *Hsf23b* expression were induced by high temperature and have similar protein localization. We also confirmed that the overexpression and knockout of *Hsf23b* as well as knockout of *Hsf23a* significantly altered heat stress tolerance, indicating that both Hsf23a and Hsf23b regulate heat stress tolerance in maize. Transcriptome analysis revealed that Hsf23b regulates a broader range of *HR* genes than Hsf23a under heat conditions. Furthermore, we demonstrated that Hsf23a physically interacts with Hsf23b in vivo, and dual-luciferase reporter assay provide evidence that Hsf23a enhanced Hsf23b’s activation activity and that promoting Hsf23b-regulated *sHSP* genes expression. Consequently, our findings highlight that *Hsf23* forms two variants by AS and both of them play crucial roles in heat stress tolerance in maize.

## Materials and Methods

### Plant materials and growth conditions

The maize inbred line B73 was maintained in our laboratory. The WT maize (KN5585) that used as the plant receptor for transgene, was provided by Weimi Biotech Co., Ltd. Maize seeds were germinated in growth chamber for 3 days at 28°C in the dark; then, maize seedlings were transplanted in pots and grown in a culture room under normal conditions with a 16 h/8 h of light/dark photoperiod, a 28°C/25°C of day/night temperature.

The Arabidopsis (*Arabidopsis thaliana*) Columbia was used as the wild-type control and the host for genetic transformation. The wild-type and transgenic plants were germinated on Murashige and Skoog basal (MS) medium for one week, and then transferred to soil and grown at 22°C in a culture room with a 16 h/8 h light/dark photoperiod.

### Identification of splice variants of *ZmHsf23*

After 42°C HS treatment for 1 h, total RNA was extracted from B73 leaves using a Total RNA Extraction Reagent (Vazyme; code R401-01-AA). The cDNA was synthesized using HiScript III RT SuperMix (Vazyme; code R323-01). The CDS of the splice variants of *ZmHsf23* were obtained by PCR. For identification of splice variants *Hsf23a* and *Hsf23b*, the cDNA of leaves with or without HS treatment (42°C and 45°C) were used. The PCR products were cloned into the pEASY-Blunt Simple Cloning Vector (TRANS; code CB111-01) and sequenced. The blast analysis of the protein sequence of Hsf23a and Hsf23b were performed to determine the splicing sites. The primers used in this study are listed in Table S1.

### Expression analysis of *Hsf23a* and *Hsf23b*

The maize inbred line B73 was used for expression analysis. The root, stem, leaf of seedlings at the three-leaf stage; and the tassel, silk and husk from maize at the V12 stage were collected for tissue-specific expression analysis.

For HS treatment, maize seedlings at the three-leaf stage were exposed directly to 42°C and sampled at the 0 h, 1 h, 3 h, 6 h, 12 h and 24 h after treatment. Total RNA was extracted from treated seedling leaves using Total RNA Extraction Reagent (Vazyme; code R401-01-AA) and cDNA synthesis was performed by reverse transcription (RT) using HiScript III RT SuperMix (Vazyme; code R323-01) according to the manufacturer’s introduction. RT-qPCR was performed using AceQ qPCR SYBR Green Master Mix (Vazyme; code Q111-02). The primer pairs for RT-qPCR were designed using the online tool of Primer3 Input 0.4.0 (https://bioinfo.ut.ee/primer3-04.0/) and listed in Table S1.

### Subcellular localization

The p1305-GFP vector (saved in our laboratory) was used for subcellular localization. The full-length CDS lacking stop codons of Hsf23a and Hsf23b were cloned into the p1305-GFP vector, constructing *35::Hsf23a-GFP* and *35::Hsf23b-GFP* fusion proteins driven by 35S promoter. The recombinant vectors were cotransformed with RFP vector (a marker for nucleus localization) into the maize protoplasts according to the previously described protocol (Cao *et al*., 2014). The p1305-GFP empty vector cotransformed with RFP vector into the protoplasts were used as a negative control. Protoplasts were cultured at multi-well plates for 18-24 h in the dark, and then the fluorescence signal were observed using a confocal laser scanning microscope (LSM710; Zeiss).

### Yeast transcriptional activity assay

The yeast strain AH109 was used for transcriptional activity assays. The full-length CDSs of Hsf23a and Hsf23b were cloned into pGBKT7 vector to generate BD-Hsf23a and BD-Hsf23b. The pGBKT7 empty vector was used as a negative control. The Hsf23b with truncated wing sequences were also cloned into the pGBKT7, resulting constructs named Hsf23b-D1 to Hsf23b-D5. All reconstructed vectors were transferred to AH109, and the transcriptional activation activity was evaluated by spot assay and X-gal staining.

### Y2H assay

The pGADT7 vector and pGBKT7 vector were used for Y2H assay. The full-length Hsf23a and Hsf23b CDSs were cloned into pGADT7 vector to generate constructs named AD-Hsf23a and AD-Hsf23b. The CDS of Hsf23b without the activation domain was cloned into pGBKT7 vector to generate construct named BD-Hsf23b-N. Different combinations of AD and BD reconstructed vectors were cotransformed into yeast AH109 cells. P53-BD + T-AD were used as positive control, BD + AD, BD + AD-Hsf23a, and BD-Hsf23b-N + AD were used as negative controls. The interaction was examined by spot assay.

### BiFC assay

The CDSs of Hsf23a and Hsf23b without the stop codons were cloned into the vectors pUC-SPYCE and pUC-SPYNE. Different combinations of cYFP and nYFP reconstructed vectors were cotransformed with RFP vector into maize protoplasts. The combinations of Hsf23a-cYFP + nYFP, cYFP + Hsf23b-nYFP were cotransformed with RFP separately as negative controls. The protoplasts isolation and transformation were performed as described previously (Cao *et al*., 2014). After transform, the protoplasts were cultured for 18-24 h in the dark, and then observed using a confocal laser scanning microscope (LSM710; Zeiss).

### Transformation and confirmation of transgenic plants

The DNA sequence encoding for the MYC tag were ligated in the 3’-end of the Hsf23a and Hsf23b CDSs lacking stop codons. The Hsf23a-MYC and Hsf23b-MYC sequences were cloned into the transformation vector pCUB, yielding *Ubi::ZmHsf23a:MYC* and *Ubi::ZmHsf23b:MYC* reconstructed vectors. The single-knockout mutant of *Hsf23a* (*hsf23a*) and double-knockout mutant of *Hsf23a* and *Hsf23b* (*hsf23a23b*) were generated by using CRISPR-Cas9. The reconstructed vector was transformed into the plant receptor WT (KN5585) maize. The overexpression and CRISPR-Cas9 transgenic plants were obtained from the Weimi Biotechnology Ltd and self-fertilized to obtain homozygotes. Total RNA was extracted from the leaves of transgenic plants at the three-leaf stage. The expression level of the transgenes were detected by RT-qPCR.

### Heat stress experiment of transgenic plants

For heat treatments of *Arabidopsis*, the WT, *OX23a* and *OX23b* seeds were germinated for 5 d and exposed to HS treatment at 45°C for 40 min, and then recovered at 22°C for 8 d before being photographed. The three-week-old seedlings of WT, *OX23a* and *OX23b* were exposed directly to 45°C for 5.5 h followed by recovery at 22°C.

For heat treatments of maize, the three-leaf stage seedlings of WT, *OE23a*, *OE23b*, *hsf23a* and *hsf23a23b* were exposed directly to 45°C HS treatment, and recovered at 28°C for 2 d. The survival rates of the plants with or without HS treatment were measured.

### TTC staining

Pollen grains from KN5585 and transgenic maize plants under control conditions or after 45°C for 1 h, were stained. Pollen grains were placed in 0.5% Triphenyltetrazolium chloride (TTC) solution and incubated at 37°C for 15 min, then the pollen viability was examined under a microscope (DM5000B, Leica, Germany).

### RNA-seq analysis

The WT, *OE23a* and *OE23b* maize seedlings that without or with 45°C HS for 1 h were used for RNA-Seq. Total RNA was extracted from the leaves of seedlings using a Total RNA Extraction Reagent (Vazyme; code R401-01-AA). Then, sequencing library construction, sequencing, and primary bioinformatics analysis were performed (GENEWIZ, Suzhou, Jiangsu Province, China). The sequencing data were analyzed as follows: Raw data were subjected to quality control using Cutadapt (v1.9.1) software. The clean data were aligned to maize reference genome via software Hisat2 (v2.0.1) and gene expression levels were estimated by the software HTSeq (v0.6.1); Differential expression analysis used the DESeq2 software, and P < 0.05 and |log_2_ Fold-Change| > 1.0 were considered as differential expressed genes (DEGs). Gene Ontology (GO) terms that annotate a list of enriched genes were identified using GOSeq (v1.34.1) with P < 0.05. Venn diagrams were plotted with online website Eveen (http://www.ehbio.com/test/venn/#/).

### Dual-luciferase assay

The CDSs of Hsf23a and Hsf23b were cloned into pGreenII 62-SK, yielding the effecter vectors. The promoter fragments of target genes were cloned into pGreenII 0800-LUC, generating the reporter vectors. Different combinations of effecter and reporter vectors were cotransformed into maize protoplasts. After incubation for 20 h, the luciferase activity (LUC/REN) was measured using the Dual Luciferase Reporter Assay Kit (Vazyme; code DL101-01).

## Results

### Alternative splicing produces two *Hsf23* transcript variants

Genome annotation showed that *ZmHsf23*, HsfA subfamily transcription factor, has two different transcript variants. The single intron is fully removed to generate the full-length transcript *Hsf23b*, while the splicing of an alternative 5’ splice site leads to a 45-nucleotide mini exon in the *Hsf23* intron, which generates a new splice transcript *Hsf23a* (Fig. 1a). To determine whether AS events occurred in *Hsf23* and these two transcript variants were present in maize, we first amplify the possible variants by reverse transcription (RT)-PCR (Table S1). Two specific bands between 1000 bp and 1500 bp were detected in maize at both normal temperature (28°C) and high temperature (42°C or 45°C), we determined that the two bands were *Hsf23a* (1098 bp) and *Hsf23b* (1053 bp), respectively (Fig. 1b). The results demonstrated that *Hsf23* underwent AS and produce *Hsf23a* and *Hsf23b* transcript variants in maize.

**Fig. 1.**
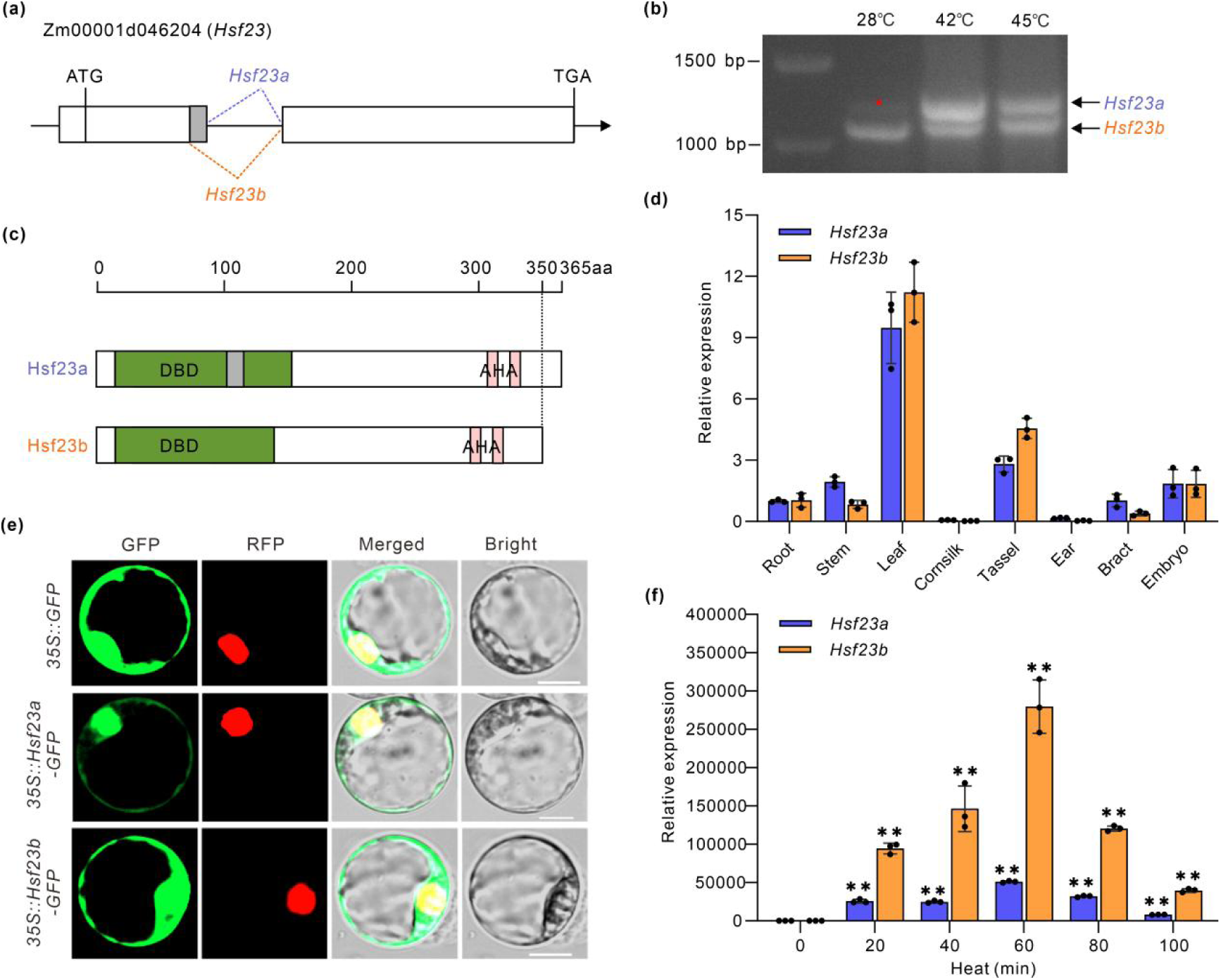
ZmHsf23 generates two transcripts and they are highly upregulated under heat stress. (a), Gene structure of *ZmHsf23* and schematic representation of two different *ZmHsf23* splicing variants. The white boxes indicate exons, the solid lines indicate introns, the dotted lines indicate splicing sites of introns, and the grey box indicates the miniexon for *Hsf23a*. (b), Detection of two *Hsf23* transcript isoforms by RT-PCR. Total RNA were extracted from maize leaves treated at 28°C or 42°C/45°C for 1 h. * indicates the splice variant *Hsf23a*. (c), Protein structures of Hsf23a and Hsf23b. DBD indicates the DNA binding domain of Hsf23a and Hsf23b. AHA indicates the activation domain of Hsf23a and Hsf23b. (d), RT-qPCR analysis of *Hsf23a* and *Hsf23b* expression in various tissues as indicated. Error bras represent means ± SD (n=3 repeats). (e), Subcellular localization of the Hsf23a-GFP and Hsf23b-GFP fusion protein in maize protoplasts. RFP is a marker for nucleus localization. The empty p1305-GFP vector served as the negative control. Bars = 10 μm. (f), Expression analysis of *Hsf23a* and *Hsf23b* under heat stress treatment. The seedlings at the three-leaf stage were treated with 42°C for 0, 20, 40, 60, 80, 100 min, and the expression levels of *Hsf23a* and *Hsf23b* were detected by RT-qPCR. Error bras represent means ± SD (n=3 repeats). Significant differences (Student’s t test: **P<0.01).

A comparison of amino acid sequences showed that Hsf23a and Hsf23b share the same modular structure, except for the DNA binding domain (DBD) (Fig. S1a). Phylogenetic analysis using the full-length protein sequences of HsfA6s revealed that Hsf23a and Hsf23b homologs are conserved in the plants examined (Fig. S1b). Hsf23 has 54% sequence similarity with AtHsfA6a and AtHsfA6b, which were positive regulators in response to salt, drought and heat stress in *Arabidopsis* (Hwang *et al*., 2014; Huang *et al*., 2016). Moreover, multiple sequence alignment demonstrated that the conservation of the DBD domain of these HsfA6 proteins, and indicated Hsf23a contained an additional wing sequence which is not normally found in the DBD of plant HSFs (Fig. S1c).

### *Hsf23a* and *Hsf23b* are strongly induced by heat stress

To investigate the potential functions of Hsf23a and Hsf23b, we examined the expression patterns of them in maize tissues and under heat-stress treatment. RT-qPCR analysis showed that both *Hsf23a* and *Hsf23b* were widely expressed in various tissues, with higher expression level in the leaf (Fig. 1d). A large-scale expression analyse showed *Hsf23* was up-regulated by HS in both B73 and CIMBL55 (Long *et al*., 2023). Not surprisingly, we noticed that *Hsf23a* and *Hsf23b* were strongly upregulated after heat treatment, furthermore, the induction of *Hsf23b* expression was much higher than that of *Hsf23a* (Fig. 1f).

To further exam the subcellular localization of Hsf23a and Hsf23b, we constructed fusion proteins containing a GFP reporter, and transiently expressed the recombinant protein in maize protoplasts. We observed that Hsf23a-GFP and Hsf23b-GFP fluorescence were distributed in the cytoplasm and nucleus (Fig. 1e), indicating that both Hsf23a and Hsf23b were cytoplasm- and nucleus-localized protein. In the localization assay, the RFP protein was used as a nucleus marker (Fig. 1e).

### Hsf23b, but not Hsf23a, enhances maize heat tolerance

Previous studies have shown that *HsfA6* genes function in the plant thermotolerance in Arabidopsis and wheat (Huang *et al*., 2016; Xue *et al*., 2015). To investigate whether Hsf23 was involved in heat responses in maize, we generated transgenic maize plants overexpressing *Hsf23a* and *Hsf23b* respectively. The Myc-tagged *Hsf23a* and *Hsf23b* were driven by the *Ubi* promoter, and the screening gene *Bar* was driven by *CaMV35S* promoter (Fig. S2a). We first examined the expression levels of *Hsf23a* and *Hsf23b* in transgenic plants. RT-qPCR analysis showed that both *Hsf23a* and *Hsf23b* expression levels were significantly increased in transgenic lines, respectively (Fig. 2a).

**Fig. 2.**
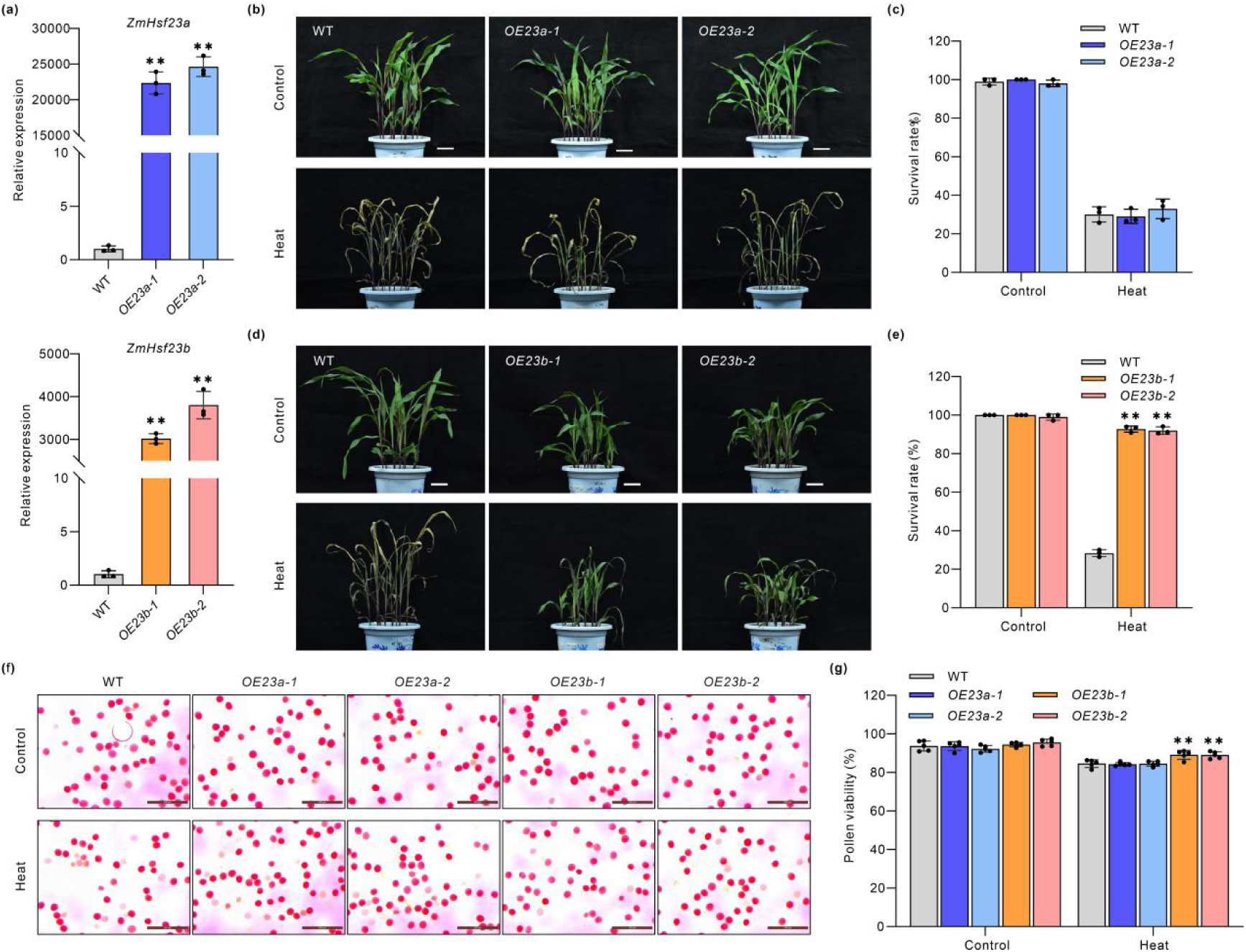
Overexpression of *Hsf23b* improve heat tolerance. (a), RT-qPCR analysis of *Hsf23a* and *Hsf23b* transcripts in the WT and transgenic plants. Bars represent means ± SD (n=3 repeats). Significant differences (Student’s t test: **P<0.01). (b), Heat phenotypes of WT and *Hsf23a* transgenic (*OE23a-1*, *-2*) seedlings. The seedlings at three-leaf stage were treated with or without 45°C for 20 h. Pictures were taken after recovery at 28°C for 2 days. Scale bars, 5 cm. (c), Survival rate of seedlings in (b). Bars represent means ± SD (n = 3 repeats). Significant differences (Student’s t test: **P < 0.01). (d), Heat phenotypes of WT and *Hsf23b* transgenic (*OE23b-1*, *-2*) seedlings under control or heat treatment (45°C, 20 h) conditions. Pictures were taken after recovery at 28°C for 2 days. Scale bars, 5 cm. (e), Survival rate of seedlings in (d). Bars represent means ± SD (n = 3 repeats). Significant differences (Student’s t test: **P < 0.01). (f), TTC staining of pollen grains. Pollen grains from WT, *Hsf23a* transgenic and *Hsf23b* transgenic lines under control conditions and after 45°C HS for 30 min, were stained. Scale bar=200 µm. (g), Relative pollen viability of pollen shown in (f). Bars represent means ± SD (n = 3 repeats). (Student’s t test: **P < 0.01).

To determine the potential functions of Hsf23 in maize, we performed heat tolerance assays in *Hsf23a* (*OE23a-1*, *2*) and *Hsf23b* (*OE23b-1*, *2*) overexpression lines. The stature of the *Hsf23a-*overexpressing plants grown at 28°C was similar to that for WT plants, however, the plant height of *Hsf23b-*overexpressing lines at 28°C was far lower than WT (Fig. S2b,c). After heat treatment, the *Hsf23b* transgenic lines exhibited a heat-tolerant phenotype (Fig. 2d) and much higher survival rates (Fig. 2e) compared with the WT plants, indicating that the overexpression of *Hsf23b* results in increased heat tolerance in maize seedlings. In contrast, no apparent difference in phenotype and survival rate were observed between WT and *Hsf23a-*overexpressing lines after heat treatment (Fig. 2b,c). ROS accumulation is an indicator of stress-induced oxidative damage, we also evaluated the accumulation of H_2_O_2_ under heat stress conditions. In agreement with their compromised heat tolerance, the H_2_O_2_ content was markedly lower in *Hsf23b-*overexpressing plants relative to the WT and *Hsf23a-*overexpressing plants (Fig. S2d). A similar results showed in transgenic *Arabidopsis* plants, the *ZmHsf23b-*overexpressing (*OX23b-1*, *2*) lines showed a heat-tolerant phenotype compared with the WT, which was accompanied by higher survival rates, whereas *ZmHsf23a-*overexpressing (*OX23a-1*, *2*) lines did not (Fig. S3).

Pollen grains are more vulnerable to heat stress compared with other tissues (Begcy *et al*., 2019). To further clarify whether Hsf23b affects the pollen viability of maize at high temperature, we performed pollen viability assay at 28°C and 45°C, and TTC was used to evaluate viability. There was no significant difference in pollen viability between WT, *Hsf23a-*overexpressing and *Hsf23b-*overexpressing plants at 28°C (control), while the values of *Hsf23b-*overexpressing plants were slightly higher than the WT after heat treatment (Fig. 2f,g). These results suggest that the Hsf23b plays a positive role in maize pollen viability under heat conditions. Taken together, these results demonstrate that overexpression of *Hsf23b*, but not *Hsf23a*, enhances the heat tolerance of maize at both the seedling and reproductive stages.

### Both *hsf23a* and *hsf23a23b* mutants are more sensitive to heat stress

To further delineate the roles of Hsf23a and Hsf23b in regulating heat tolerance, we generated *hsf23a* single-knockout mutant and *hsf23a23b* double-knockout mutant by using CRISPR-Cas9. The *hsf23a-1* mutant harbored a 1-bp insertion and *hsf23a-2* carried a 1-bp deletion in *Hsf23a* (Fig. 3a), leading to frame shifts in the open-reading frame and premature termination of translation (Fig. S4a). In addition, the *hsf23a23b-1* mutant and *hsf23a23b-2* contained 38-bp and 37-bp deletion respectively, leading to the loss of *Hsf23* function (Fig. 3b, S4a). After selection and identification, we obtained homozygous *hsf23a* and *hsf23a23b* mutant lines without the Cas9 transgene for further study.

**Fig. 3.**
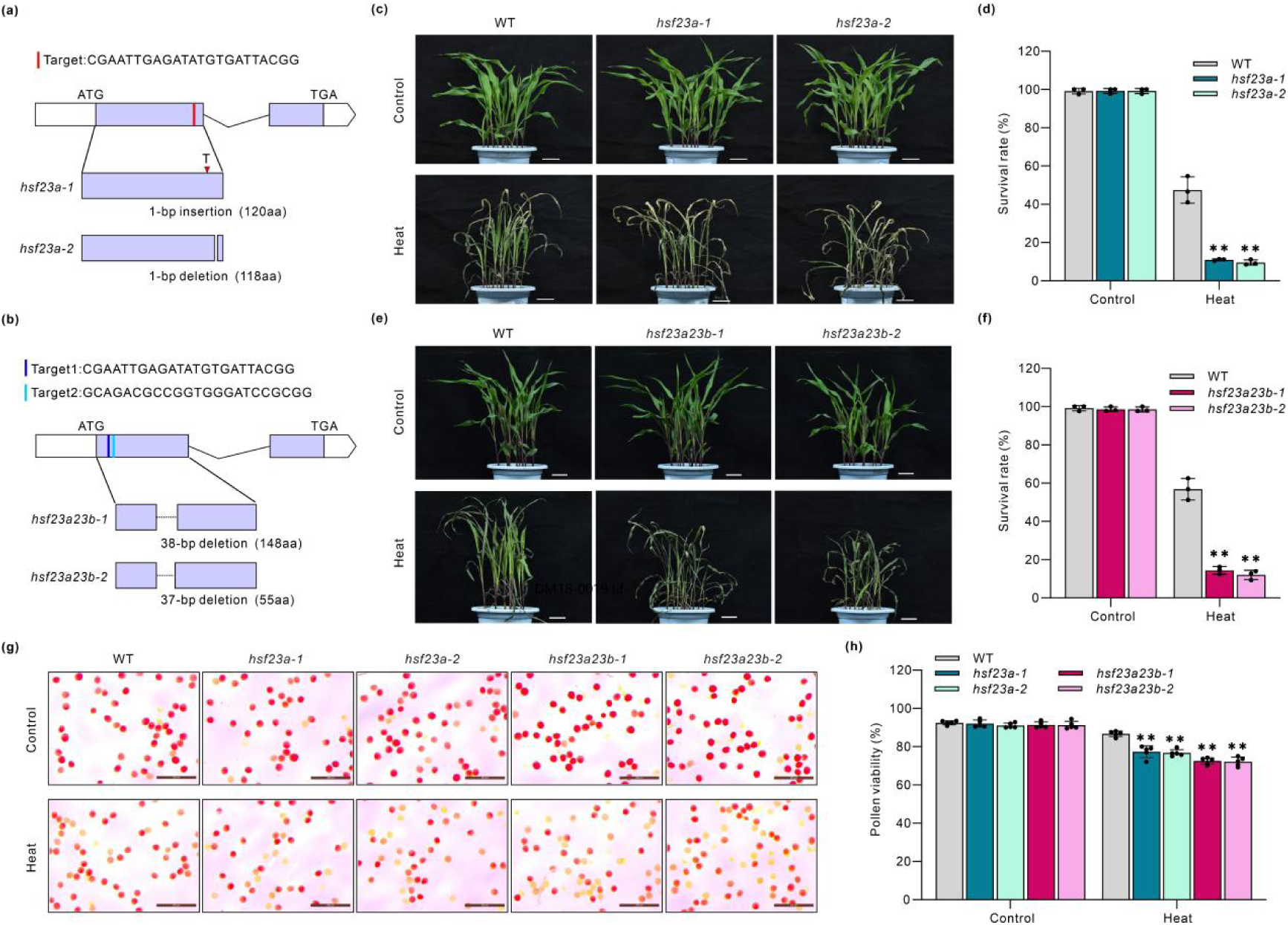
Knockout mutants of *hsf23a* and *hsf23a23b* are more sensitive to heat stress. (a, b), Schematic diagram of the *hsf23a* mutants (a, *hsf23a-1*,*2*) and *hsf23a23b* mutants (b, *hsf23a23b-1*,*2*) generated by CRISPR/Cas9-mediated genome editing. (c), Heat phenotypes of WT and *hsf23a* mutant seedlings. The seedlings at three-leaf stage were treated with or without 45°C for 14 h. Pictures were taken after recovery at 28°C for 2 days. Scale bars, 5 cm. (d), Survival rate of seedlings in (c). Bars represent means ± SD (n = 3 repeats). Significant differences (Student’s t test: **P < 0.01). (e), Heat phenotypes of WT and *hsf23a23b* mutant seedlings under control or heat treatment (45°C, 14 h) conditions. Pictures were taken after recovery at 28°C for 2 days. Scale bars, 5 cm. (f), Survival rate of seedlings in (e). Bars represent means ± SD (n = 3 repeats). Significant differences (Student’s t test: **P < 0.01). (g), TTC staining of pollen grains. Pollen grains from WT, *hsf23a* mutants and *hsf23a23b* mutants under control conditions or after 45°C for 30 min, were stained. Scale bar=200 µm. (h), Relative pollen viability of WT, *hsf23a* mutants and *hsf23a23b* mutants before and after heat treatments. Bars represent means ± SD (n = 3 repeats). (Student’s t test: **P < 0.01).

Under normal conditions, there was no significant difference in plant height between *hsf23*, *hsf23a23b* mutants and WT plants (Fig. S4b,c). Unexpectedly, in contrast to *Hsf23a*-overexpressing lines, *hsf23a-1* and *hsf23a-2* displayed a heat-sensitive phenotype in comparison with the WT under heat conditions (Fig. 3c). Consistent with this observation, the survival rates were significantly lower and H_2_O_2_ contents were higher in *hsf23a* mutants relative to the WT after heat treatment (Fig. 3d, S4d), indicating that Hsf23a is involved in regulating heat tolerance in maize. We also observed heat-sensitive phenotype (Fig. 3e), a lower survival rate (Fig. 3f) and much higher H_2_O_2_ accumulation (Fig. S4d) for *hsf23a23b* mutants under heat conditions. Furthermore, the pollen viability of knockout mutants was similar to that of WT under control conditions, but the values of these mutants dramatically decreased after heat treatment (Fig. 3g,h). These results indicate that both *hsf23* and *hsf23a23b* mutants were highly susceptible to heat stress.

### Hsf23b affects the broader heat-responsive genes transcription than Hsf23a during heat stress

To investigate the roles of Hsf23a and Hsf23b in regulating heat stress tolerance, we performed RNA-seq analysis using the leaves of WT, *Hsf23a*-overexpressing (*OE23a*) and *Hsf23b*-overexpressing (*OE23b*) seedings grown at 28°C or exposed to 45°C heat treatment for 1 h. Thousands of differentially expressed genes (DEGs) in response to HS were identified in *OE23a*, *OE23b* and WT plants (Fig. S5, Table S2). We identified 5415 DEGs, consisting of 2552 downregulated and 2863 upregulated genes, in WT at HS conditions (45°C) compared with normal conditions (28°C). In *OE23a* plants, there were far more up-regulated genes than down-regulated genes at 45°C compared with 28°C. However, down-regulated genes outnumbered up-regulated genes in the *OE23b* plants when comparing 45°C to 28°C (Fig. S5a-c).

We also identified DEGs in the comparison between *OE23a*, *OE23b* and WT at normal and HS conditions (Table S3). In normal conditions, we identified 362 DEGs in *OE23a* and 2689 DEGs in *OE23b* relative to WT, respectively. There were 1405 up-regulated DEGs in *OE23b* compared with WT at normal conditions (Fig. 4a, S5e), while only 323 up-regulated genes in *OE23a* (Fig. 4a, S5d). Gene ontology (GO) analysis revealed that these DEGs up-regulated in *OE23b* were mainly enriched for terms related to “response to heat,” “response to stress,” and “response to hydrogen peroxide” (Fig. S6c), while the up-regulated DEGs in *OE23a* at normal conditions were not specific enriched for stress response functions (Fig. S6a). We also found that the DEGs involved in inositol biosynthetic process, photosynthesis, carbohydrate biosynthetic process, and gibberellin metabolic process were down-regulated in *OE23b* (Fig. S6d). This result is consistent with the observation that increase of *ZmHsf23b* expression level leads to dwarf phenotype under normal conditions (Fig. S2b, S6d).

**Fig. 4.**
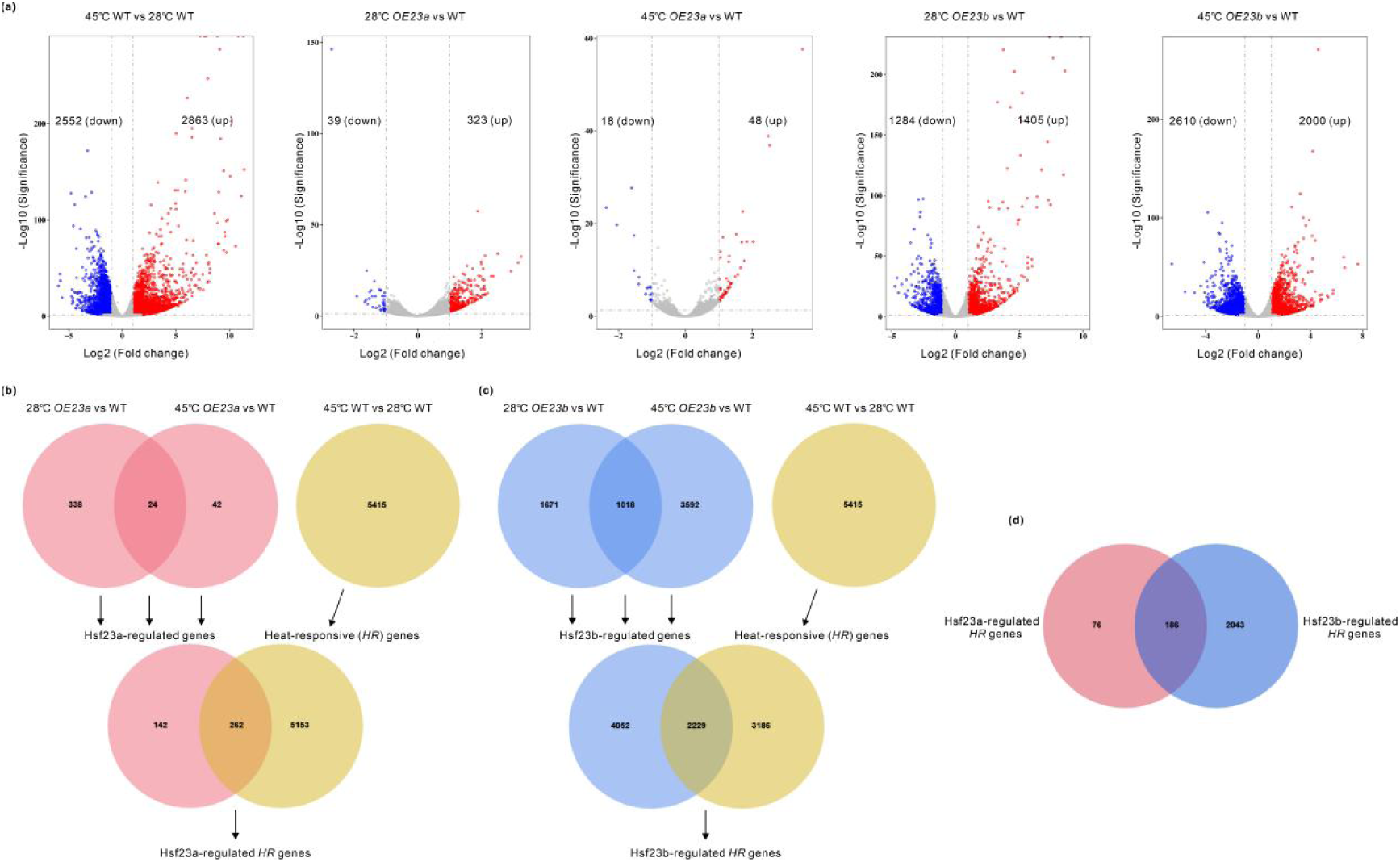
Transcriptome analysis of Hsf23a-regulated and Hsf23b-regulated *HR* genes. (a), Volcano plots showing genes upregulated or downregulated in the comparison between *OE23a*, *OE23b* and WT in response to HS. (b), Venn diagrams showing Hsf23a-regulated DEGs (top, left), heat-regulated DEGs (top, right), and both heat- and Hsf23a-regulated DEGs (bottom). (c), Venn diagrams showing Hsf23b-regulated DEGs (top, left), heat-regulated DEGs (top, right), and both heat- and Hsf23b-regulated DEGs (bottom). (d), Venn diagram representing the overlap of Hsf23a- and Hsf23b-regulated *HR* genes.

Given the finding that Hsf23a and Hsf23b play different as well as similar roles in heat stress response, we hypothesized that Hsf23a and Hsf23b might regulate different downstream genes in maize. To test this hypothesis, we identified Hsf23a or Hsf23b-regulated DEGs by comparing the RNA-seq data from the *OE* plants and WT at 28°C or 45°C. A total of 5415 genes were defined as heat-responsive (*HR*) genes in WT plants with heat treatment compared with plants without heat treatment (Fig. 4a). We identified 404 Hsf23a-regulated DEGs, including 362 DEGs from *OE23a* versus WT at 28°C and 66 from *OE23a* versus WT at 45°C (Fig. 4a,b). By comparing the 5415 and 404 DEGs, we obtained 262 overlapping genes that are regulated by both heat and Hsf23a (Fig. 4b, Table S4). In the same way, we also identified 2229 *HR* genes regulated by Hsf23b, which accounts for 41% of *HR* genes and 35% of Hsf23b-regulated genes (Fig. 4c, Table S4). Moreover, we compared the 262 Hsf23a-regulated *HR* genes with 2229 Hsf23b-regulated *HR* genes, and found that a set of 186 common *HR* genes were regulated by both Hsf23a and Hsf23b, while 76 and 2043 *HR* genes were only regulated by Hsf23a and Hsf23b, respectively (Fig. 4d). Therefore, these results demonstrate that Hsf23b regulates a broader range of *HR* genes than Hsf23a in heat stress response.

### Hsf23a and Hsf23b modulate heat stress response through different downstream targets

RNA-seq data showed that 48 and 2000 genes were upregulated by heat in *OE23a* and *OE23b* compared with the WT, respectively (Table S2). As *Hsf23a* and *Hsf23b* positively regulated heat stress response, we further focused on 11 and 371 upregulated genes under heat stress conditions, which showed higher expression in *OE23a* or *OE23b* plants compared with the WT at 45°C (Fig. 5a,b, Table S5). We performed gene ontology (GO) analysis of these 371 genes to explore the biological processes involving Hsf23b. The enrichment analysis indicated that these DEGs are significantly enriched for terms such as “protein folding”, “response to abiotic stimulus”, and “response to stress” (Fig. 5c).

**Fig. 5.**
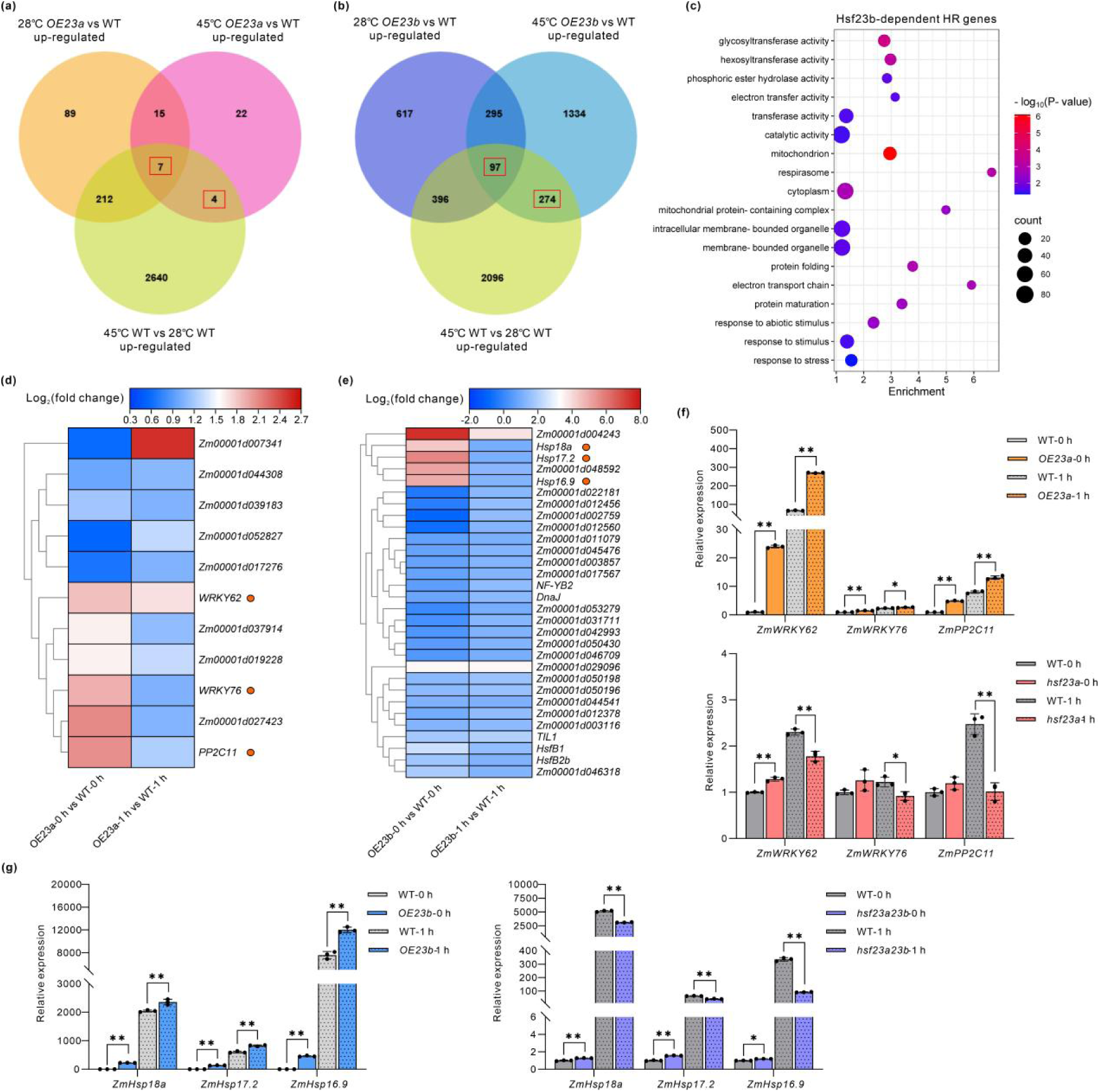
Hsf23a and Hsf23b positively regulate *HR* genes under heat stress conditions. (a), Venn diagram showing the differentially upregulated genes in *OE23a* and WT plants. (b), Venn diagram showing the differentially upregulated genes in *OE23b* and WT plants. (c), GO analysis showing enrichment terms of Hsf23b-dependent *HR* genes under heat stress in (b). In total, 371 *HR* genes that were upregulated in *OE23b* under heat stress conditions (fold change > 1, p-value < 0.05) compared with WT were used for the GO analysis. (d), Heatmap of expression patterns of Hsf23a-dependent *HR* genes as shown in (a). Heatmap shows the Log2 (TPM) values of 11 DEGs. (e), Heatmap showing the expression patterns of enriched genes in the GO term Protein folding, Response to stress shown in (c). Heatmap shows the Log2 (TPM) values of 30 DEGs. (f), RT-qPCR analysis of *PP2C11*, *WRKY62* and *WRKY76* expression in WT, *OE23a*, and *hsf23a* seedlings before and after heat treatment. Bars represent means ± SD (n=3 repeats). Significant differences (Student’s t test: **P<0.01, *P<0.05). (g), RT-qPCR analysis of *Hsp16.9*, *Hsp17.2* and *Hsp18a* expression in WT, *OE23b*, and *hsf23a23b* seedlings before and after heat treatment. Bars represent means ± SD (n=3 repeats). Significant differences (Student’s t test: *P<0.05, **P<0.01).

To investigate the potential target genes underlying Hsf23a- and Hsf23b-regulated heat stress tolerance, we analyzed the 11 Hsf23a-regulated DEGs by constructing heatmaps as well as 30 Hsf23b-regulated DEGs enriched in protein folding and response to stress. Among the 11 Hsf23a-dependent *HR* genes, three of them encode transcription factors, including PP2C11, WRKY62 and WRKY76 (Fig. 5d). *WRKY* genes encode transcription factors that play multiple roles in plant responses to abiotic stress, including heat stress (Ashraf *et al*., 2018). The type-2C protein phosphatases (PP2Cs) are the core components of ABA signaling, and these proteins are involved in the response to abiotic stress through ABA-dependent pathway (Wei *et al*., 2014; Miao *et al*., 2020). The heatmaps revealed that these three genes showed higher expression in *OE23a* compared with the WT under both normal and HS conditions (Fig. 5d). These results suggest that Hsf23a positively regulates heat stress tolerance by modulating the expression of *PP2C11*, *WRKY62* and *WRKY76.* Notably, the expression of *sHSP* genes in *OE23b*, including *Hsp16.9, Hsp17.2,* and *Hsp18a,* was highly up-regulated relative to WT before heat treatment, and this up-regulation also pronounced after heat treatment (Fig. 5e). The up-regulation of *HSPs* via an HSFs-dependent system is the central part of heat stress responses (McLoughlin *et al*., 2016). Therefore, we conjectured that the effects of Hsf23b during the heat stress response might associated with *sHSP* gene expression. Moreover, we designed specific primers for these six genes and performed RT-qPCR analysis. *PP2C11*, *WRKY62* and *WRKY76* transcript levels were upregulated in the *OE23a* plants and were downregulated in *hsf23a* mutants compared with the WT after heat treatment (Fig. 5f). After heat treatment, three *sHSP* genes showed significantly higher expression in the *Hsf23b* overexpression lines and lower expression levels in *hsf23a23b* mutants than in the WT (Fig. 5g). These results support the notion that Hsf23a and Hsf23b regulate heat stress tolerance through target different downstream *HR* genes.

To ascertain whether Hsf23a or Hsf23b directly activated the transcription of these potential targets, we performed Dual-LUC reporter assays. The promoter of target genes (1.5 kb upstream from the start codon) were fused with the luciferase coding sequence, forming the reporter vectors; the coding sequence of Hsf23a or Hsf23b were driven by the CaMV 35S promoter, yielding the effector vectors (Fig. 6a,c). When PP2C11 pro-LUC, WRKY62 pro-LUC or WRKY76 pro-LUC was individually transfected into tobacco leaves, basal LUC fluorescence signals were detected. When them were cotransfected with 35S-Hsf23a respectively, the tobacco leaves showed significantly higher fluorescence signals, indicating that activation of the *PP2C11*, *WRKY62* and *WRKY76* promoters by Hsf23a (Fig. 6b). Similarly, strong LUC signals were detected in the leaf regions co-expressing the 35S-Hsf23b effector and sHsp pro-LUC reporters, confirming Hsf23b activates the promoter activity of *Hsp16.9, Hsp17.2,* and *Hsp18a* (Fig. 6d). We also performed dual-luciferase transcriptional activity assays in maize protoplasts. Different combinations of the reporter and effector vectors were co-transformed into the maize protoplasts, and the relative luciferase activity was measured (Fig. 6e,f). Consistent with the qualitative test shown in Fig. 6b, Hsf23a showed significant activation on *PP2C11*, *WRKY62* and *WRKY76* promoters in this assay (Fig. 6e). As expected, Hsf23b showed significant activation on *Hsp16.9, Hsp17.2,* and *Hsp18a* promoters (Fig. 6f). These results provide evidence that Hsf23a and Hsf23b target different genes during heat stress responses.

**Fig. 6.**
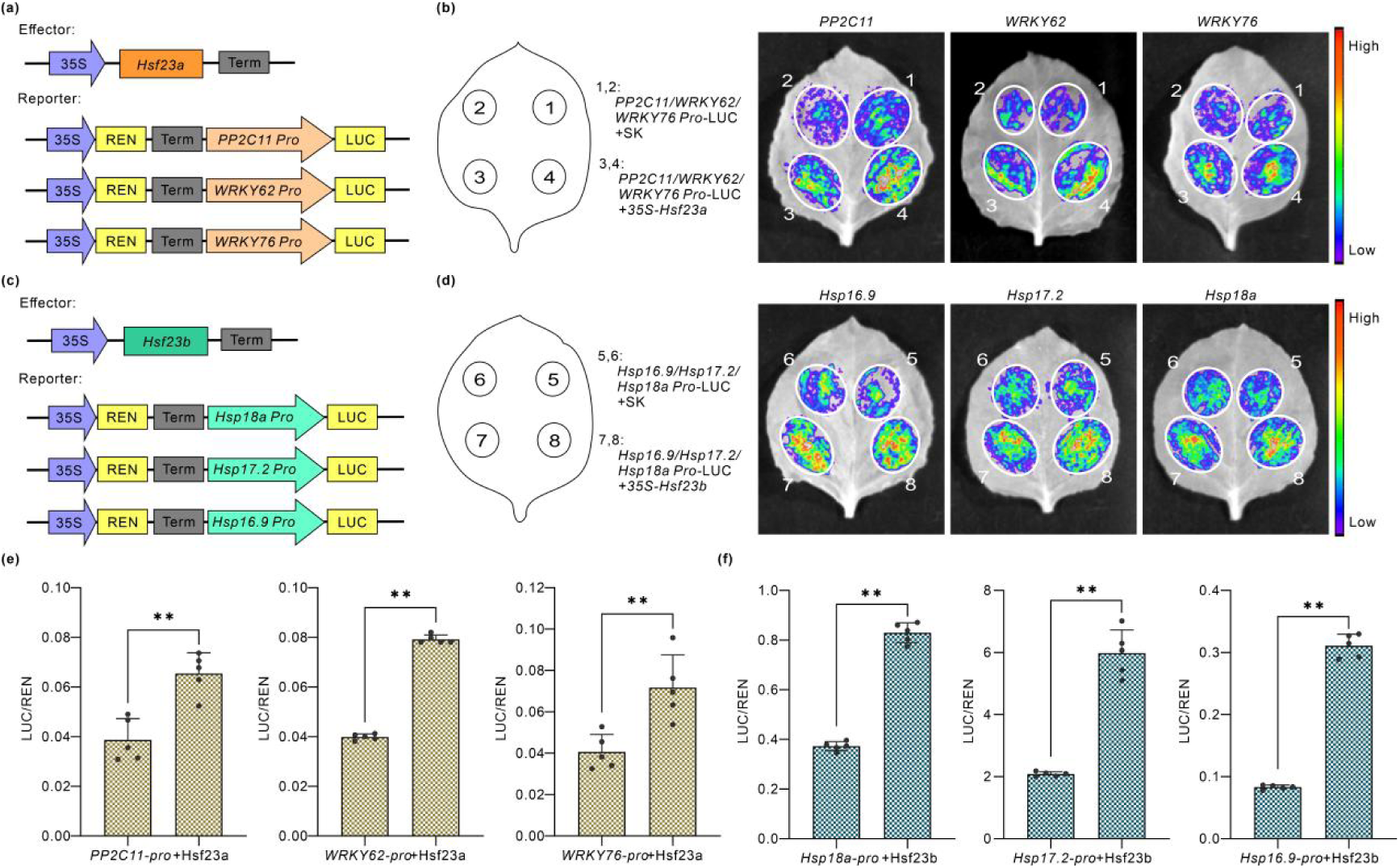
Hsf23a and Hsf23b modulate heat stress response through different downstream targets. (a), The effector and reporter constructions used for the *PP2C11*, *WRKY62* and *WRKY76* promoters transactivation assay. (b), Analysis of the transcriptional activation properties of Hsf23a in tobacco leaves. Different combinations were co-transformed in tobacco leaves. (c), The effector and reporter constructions used for the *sHSP* promoters transactivation assay. (d), Analysis of the transcriptional activation properties of Hsf23b in tobacco leaves. Different combinations were co-transformed in tobacco leaves. (e), Dual-luciferase reporter assay shows the direct regulation of Hsf23a on the transcriptional activation of *PP2C11*, *WRKY62* and *WRKY76*. Different combination of vectors (a) was expressed in maize protoplasts. Error bars represent SE (n = 5). Significant differences (Student’s t test: **P < 0.01). (f), Dual-luciferase reporter assay shows the direct regulation of Hsf23b on the transcriptional activation of *Hsp16.9*, *Hsp17.2* and *Hsp18a*. Different combination of vectors (c) was expressed in maize protoplasts. Error bars represent SE (n = 5). Significant differences (Student’s t test: **P < 0.01).

### Hsf23a physically interacts with Hsf23b

The protein interactions between HSFs are play important roles in the regulation of gene expression (Friedrich *et al*., 2021; Wu *et al*., 2024). To determine whether Hsf23a and Hsf23b formed homologous or heterologous interactions, we perform a yeast two-hybrid (Y2H) assay. Since Hsf23a and Hsf23b contain C-terminal activation domain, we first detect the self-activation activity of them (Fig. S7). It was observed that Hsf23b has transcriptional activation activity in yeast cells and its activation domain lied between amino acids 291 and 350. In contrast, Hsf23a has no transcriptional activation activity due to the insertion of wing sequence in the DBD domain (Fig. S7). Then, we used the Hsf23b-N protein, which lack of activation domain, for the Y2H assay. Yeast cells transformed with Hsf23a/Hsf23a, Hsf23b/Hsf23b-N, and Hsf23a/Hsf23b-N combinations grew well on the deficient medium, indicating that interactions of Hsf23a with itself, Hsf23b with itself, and Hsf23a with Hsf23b (Fig. 7a). Moreover, we used a BiFC assay to confirm and localize the interactions. Co-expression of Hsf23a-cYFP and Hsf23b-nYFP in maize protoplasts resulted in YFP signals in the nucleus, demonstrating that Hsf23a interacted with Hsf23b occur in the nucleus (Fig. 7b). The interaction between Hsf23a and Hsf23b was further confirmed by LCI assays. The luciferase luminescence were observed in the combination of Hsf23a/Hsf23a and Hsf23b/Hsf23b homologous interactions and Hsf23a/Hsf23b heterologous interaction, confirming a strong interaction between Hsf23a and Hsf23b (Fig. 7c). Together, these results demonstrate that Hsf23a can interact with Hsf23b to form a heterodimer.

**Fig. 7.**
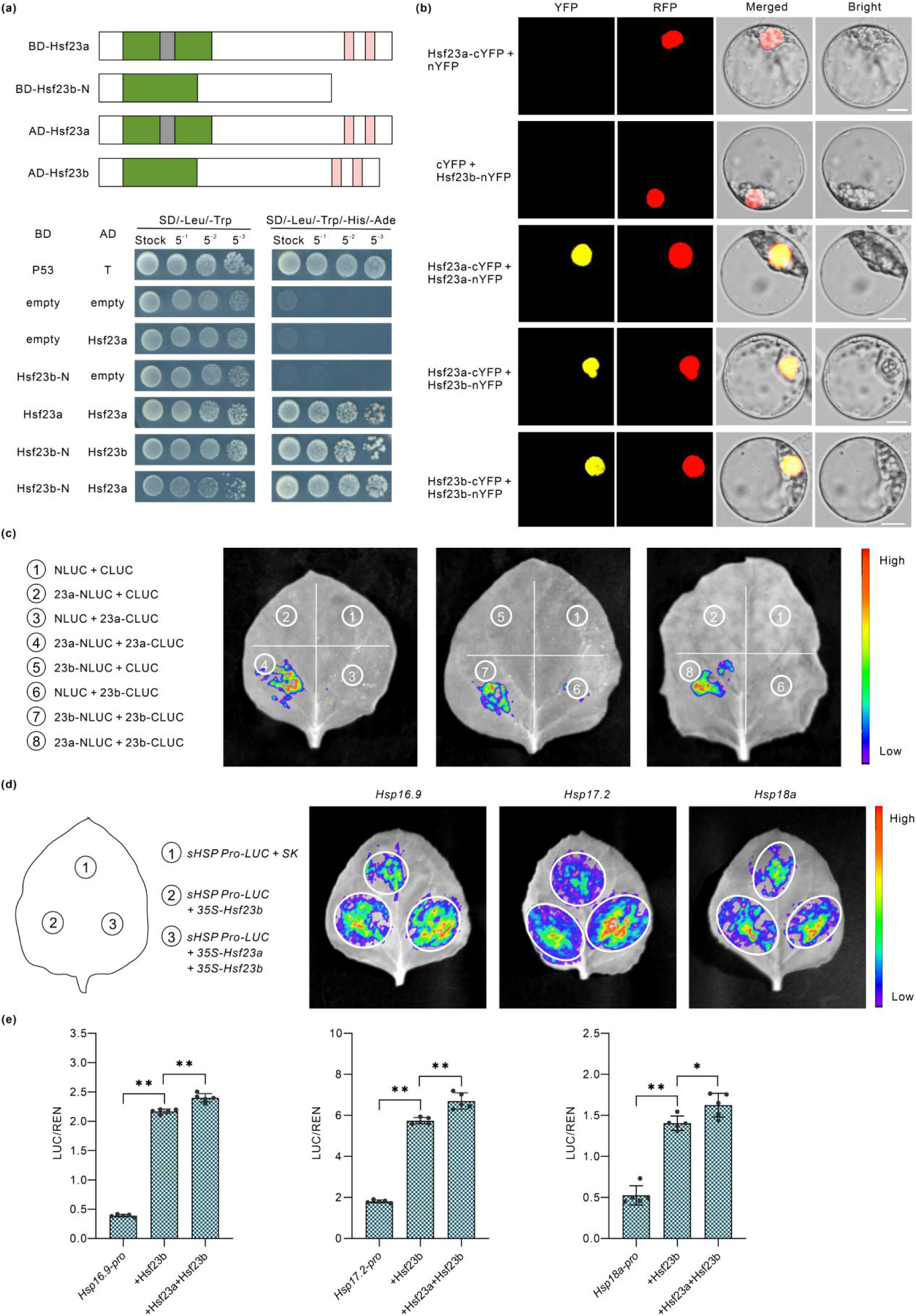
Hsf23a interacts with Hsf23b to promotes Hsf23b-dependent *sHSP* expression. (a), Interaction of Hsf23a and Hsf23b in Y2H assay. AD, GAL4 activation domain; BD, GAL4 DNA binding domain; Protein–protein interactions were examined by yeast cell growth on the SD/-Leu/-Trp/-His/-Ade plate. (b), Interaction of Hsf23a and Hsf23b in BiFC assay. Different combinations of the nYFP and cYFP fusion constructs were co-transformed into maize protoplasts, and the BiFC signals were analyzed using a fluorescence microscope. RFP is a marker for nucleus localization. Scale bars, 10 µm. (c), Luciferase complementation imaging (LCI) assay of Hsf23a and Hsf23b. Different combinations of the CLUC and NLUC fusion constructs were transiently coexpressed in *Nicotiana benthamiana* leaves. The fluorescent signal represents protein-protein interaction. (d), Transactivation of the *sHSP* promoters by Hsf23a and Hsf23b in *Nicotiana benthamiana* leaves. (e), Dual-LUC reporter assay showing that Hsf23a positively regulates Hsf23b-dependent *Hsp16.9*, *Hsp17.2* and *Hsp18c* transcriptions in maize protoplasts. Error bars represent SE (n = 5). Significant differences (Student’s t test: *P < 0.05, **P < 0.01).

### Hsf23a promotes Hsf23b-regulated expression of *sHSP* genes involved in heat stress response

Given that Hsf23a interacts with Hsf23b, we tested whether Hsf23a mediate the expression of *sHSP* genes which were regulated by Hsf23b. We co-expressed Hsf23a and Hsf23b in a dual-luciferase transient transcriptional activity assay in tobacco leaves (Fig. 7d). Consistent with earlier findings, the strong LUC signals in the leaf regions expressing Hsf23b and different *sHSP* promoters confirmed Hsf23b transactivates the *Hsp16.9, Hsp17.2,* and *Hsp18a* promoters (Fig. 6d). When *Hsf23a* and *Hsf23b* were co-expressed, the leaf regions showed higher LUC signal than the expression of Hsf23b alone, revealing Hsf23a enhanced Hsf23b’s activation of the *sHSP* promoters (Fig. 7d). Then, we performed dual-luciferase reporter assay in maize protoplasts, and the luciferase activity (LUC/REN) was measured (Fig. 7e). It was revealed that Hsf23b alone showed significant activation on *sHSP* promoters; in addition, the activity was further enhanced when *Hsf23a* was co-expressed with *Hsf23b* (Fig. 7e). Overall, these results suggest that Hsf23a interacts with Hsf23b to promotes Hsf23b’s activation of the *sHSP* genes expression.

## Discussion

Heat stress severely affects crop yield and quality, as high temperatures reduce photosynthesis and energy metabolism, and disrupt flower formation and pollination processes (Lohani *et al*., 2020; Moore *et al*., 2021). Therefore, there is an urgent need to uncover the mechanisms underlying heat stress tolerance in plant to develop crop cultivars with improved thermotolerance. HSFs play vital roles in regulating plant HSR (Akerfelt *et al*., 2010; Ohama *et al*., 2017), however, the molecular mechanism of HSFs in heat stress tolerance in maize have been unclear. Here, we demonstrated that the AS of *Hsf23* is an important adaptative mechanism that positively regulates heat stress tolerance in maize by modulating the expression of a set of *HR* genes.

The expression and activation of HSFs are strictly regulated at different levels, and HS-induced AS is an important post-transcriptional regulation of plant *HSFs* (Meng *et al*., 2016; Ohama *et al*., 2017). The AS induced by HS is observed for *Arabidopsis HSFA2*, *A7b*, *A4c* and *B1*, grape *HSFA2*, rice *HSFA2d*, and lily *HSFA3s*, and all of their splice variants are produced by intron retention under HS conditions (Liu *et al*., 2013; Cheng *et al*., 2015; Jiang *et al*., 2017; Wu *et al*., 2019). In this study, we found the AS of *Hsf23* generates two transcript variants. The fully remove of intron generates *Hsf23b* transcript, while the splicing of 5’ splice site in intron leads to a cryptic miniexon in *Hsf23a* transcript (Fig. 1a). A study in *Arabidopsis* showed that *AtHsfA2* generated a splice variant *AtHsfA2-III* through a cryptic 5’ splice site in the intron under severe HS, the splice mode of *AtHsfA2-III* is somewhat similar to that of *Hsf23a*; However, AtHsfA2-III contained a premature stop codon (PTC) and encoded a truncated AtHSFA2 protein, whereas Hsf23a contained an additional wing sequence in DBD and encoded a larger Hsf23 protein (Fig. 1c); the study also reported HS-induced intron retention and PTC also occurs in *HsfA4c*, *HsfA7b*, *HsfB1*, and *HsfB2a* (Liu *et al*., 2013). Therefore, severe HS-induced AS is the most frequent event among plant *HSF* genes, and the splice mode of plant *HSFs* is diversified.

Hsf23 has a typical structure of class A, featuring the AHA motifs which mainly consist of acidic and hydrophobic amino acids. The HsfAs have one or more AHA motifs located in their C-terminal activation domain, which mediate HsfAs function as transcription activators (Meng *et al*., 2016). Here, we observed that Hsf23b has two AHA motifs in its C-terminal sequence and showed transactivation activity in yeast cells (Fig. S7), which is consistent with the reported conclusions. Interestingly, even though Hsf23a also contain AHA motifs, it had no transactivation activity in yeast cells, and this might be explained by the repression role of the additional wing sequences in DBD domain (Fig. S7). The yeast HSF belongs to the winged helix protein family, and almost of them contain the wing sequence in the DNA-binding domain, and deletion of the wing sequence suppress the transcriptional activity of HSFs (Cicero *et al*., 2001). In this study, our transactivation assays showed that insertion of wing sequence in Hsf23b DBD lead to the lack of transactivation activity. Therefore, having AHA motifs does not necessarily mean that the HSF has transactivation activity.

HSFs are the conserved and central regulators of the HSR in eukaryotic. Among plant *HSFs*, *HsfAs*, especially *HSFA1* and *A2*, are the dominant transcriptional activators of HS-regulated gene expression and are indispensable for evoking the HSR in plants (Li *et al*., 2018; Andrasi *et al*., 2021). In our study, *Hsf23a* and *Hsf23b* expression was strongly induced by heat treatment (Fig. 1f), implying that *Hsf23a* and *Hsf23b* might have roles in maize heat stress tolerance. Overexpression of *AtHSFA6b* or *CaHSFA6a* improve the thermotolerance of the transgenics (Huang *et al*., 2016; Mou *et al*., 2024). Our study found that overexpression of *Hsf23b* enhanced heat stress tolerance in transgenic plants (Fig. 2), while knockout it caused the opposite effect (Fig. 3), implying that Hsf23b was a positive regulator of plant heat stress tolerance. In addition, we also observed that *Hsf23a* overexpression lines were no difference in heat tolerant compared with WT (Fig. 2), but the *hsf23a* mutants were more sensitive to heat (Fig. 3), suggesting that the loss of Hsf23a function results in decreased heat stress tolerance. These results indicated that both Hsf23a and Hsf23b were required for heat stress tolerance in maize, but there might exist different mechanisms of Hsf23a and Hsf23b.

HSFAs function as central activators of the HSR, they are known to bind to the heat stress elements (HSEs) in the target promoters, and subsequently activates the downstream heat-responsive gene expression (Xue *et al*., 2015; Ohama *et al*., 2016). We demonstrated that Hsf23a upregulated 11 *HR* genes under heat stress conditions, while Hsf23b upregulated 371 *HR* genes (Fig. 5), perhaps due to the differences in transcription activation function or gene regulation pathways of these isoforms. It is speculated that the appear of wing sequence in DBD domain affects the DNA-binding properties of Hsf23a and Hsf23b. HSF binds DNA as a trimer, and additional trimers can bind DNA co-operatively. The ‘wing’ in the HSF DBD does not appear to contact DNA. A previously study indicates that the wing is involved in protein-protein interactions and is likely affects the DNA-binding specificity and affinity of the HSF trimers (Cicero *et al*., 2001). We speculated that Hsf23a may occupy the promoter HSE position with no or weaker transcription ability but cooperates with other factors for transactivation to exhibit activation capability. Similarly, HSFBs and HSFCs have not AHA motifs in their C-terminus, but the reports of tomato SlHSFB1 and pepper CaHSFB2a have shown that they can activate the target genes by binding the promoters (Ashraf *et al*., 2018; Fragkostefanakis *et al*., 2019). All in all, the specific basis needs further research. WRKY TFs are one of the largest TF families in higher plants and play critical roles modulating diverse signaling pathways during stress responses (Chen *et al*., 2010; Ashraf *et al*., 2018). In pepper, CaHsfB2a positively regulates plant tolerance to high-temperature by forming transcriptional cascades and positive feedback loops with CaWRKY6 and CaWRKY40 (Ashraf *et al*., 2018). Moreover, WRKYs were also found to be involved in regulation of the ABA-dependent stress responses. For example, *Arabidopsis* WRKY40 was shown to enhance abscisic acid (ABA)-mediated salt and osmotic stress tolerance by antagonizing WRKY18 and WRKY60 functions (Chen *et al*., 2010). Zm*WRKY82* and *ZmWRKY83* genes (called *WRKY76* and *WRKY62* in this study), the orthologs of *AtWRKY40*, were might involved in the transcriptional regulation of ABFs/AREBs (Hu *et al*., 2021). Here, we confirmed that Hsf23a directly binds to the promoters of *WRKY62* and *WRKY76*, promoting their expressions along with other downstream genes (Fig. 6). In addition, our results showed that Hsf23a directly regulates *PP2C11* transcription (Fig. 6). Group A PP2Cs were reported as negative regulators in ABA signaling pathway and acted as key regulators of stress responses (Komatsu *et al*., 2013; Miao *et al*., 2020). *PP2C11* (published *ZmPP112*), the homolog of rice *OsPP2C1*, was highly induced in response to abiotic stresses (Wei *et al*., 2014). These data suggested Hsf23a might contribute to heat stress tolerance through ABA-mediated heat responses. Consistent with previous studies, we determined that Hsf23b binds to and upregulates *sHSPs* in response to heat stress (Fig. 6). HSF-HSP network is the major component of heat stress responses. As molecular chaperones, HSPs contribute to protein homeostasis in plants, supporting protein refolding and misfolded protein degradation in response to heat stress (McLoughlin *et al*., 2016). Heterologous expression of wheat *sHSP17.4* and *sHSP25.9* in *Arabidopsis* enhanced plant thermotolerance (Feng *et al*., 2019; Wang *et al*., 2023). A recent study found that TaHSFA6e-TaHSP70 signaling module coordinately regulates heat response in wheat (Wen *et al*., 2023). Taken together, our results indicated that Hsf23a and Hsf23b might modulate heat stress tolerance in maize via different signaling pathway.

HSFs can often form homologous or heterologous oligomers to participate in the regulation of gene expression, and such interactions often occur between same HSF class. In tomato, HsfA1 interacts with HsfA2 to form heterologous complexes, resulting in strong synergistic activation of downstream HS-related gene expression (Chan-Schaminet *et al*., 2009). AtHSFA2 interacts with AtHSFA3 to efficiently promote the transcriptional activation of memory-related genes during the HS memory in *Arabidopsis* (Friedrich *et al*., 2021). Our results showed that Hsf23a interacted with itself, Hsf23b interacted with itself, as well as Hsf23a interacted with Hsf23b (Fig. 7). More importantly, we found co-expression of Hsf23a promoted Hsf23b to activate its target genes (Fig. 7), suggesting that Hsf23a and Hsf23b formed a complex with stronger transactivation ability, and further increased the expression of *sHSPs* genes.

In summary, our findings demonstrated that the AS of *Hsf23* play crucial roles in regulating heat stress tolerance in maize, since *Hsf23* could generate two protein isoforms that play different roles under heat conditions. According to our findings, we propose a working model by which Hsf23 regulates plant heat stress tolerance. Under heat stress conditions, the enhancement of *Hsf23* transcription and the AS of *Hsf23* increases *Hsf23a* and *Hsf23b* transcript levels. The increase in *Hsf23a* expression leads to the activation of ABA signaling genes to promote ABA-mediated heat responses, and higher *Hsf23b* transcript levels under heat stress enhances the activation of *sHSP* genes, which modulates plant thermotolerance. Simultaneously, Hsf23a interacts with Hsf23b to greatly enhance the transcriptional activation activity of Hsf23b toward its target genes (Fig. 8). Therefore, the AS of *Hsf23* produces two variants that co-operate with each other to help plants adapt to heat stress. Further investigation of the molecular mechanism mediated by the AS of *Hsf23* in the heat stress response will contribute to breed heat-tolerant crops in the future.

**Fig. 8.**
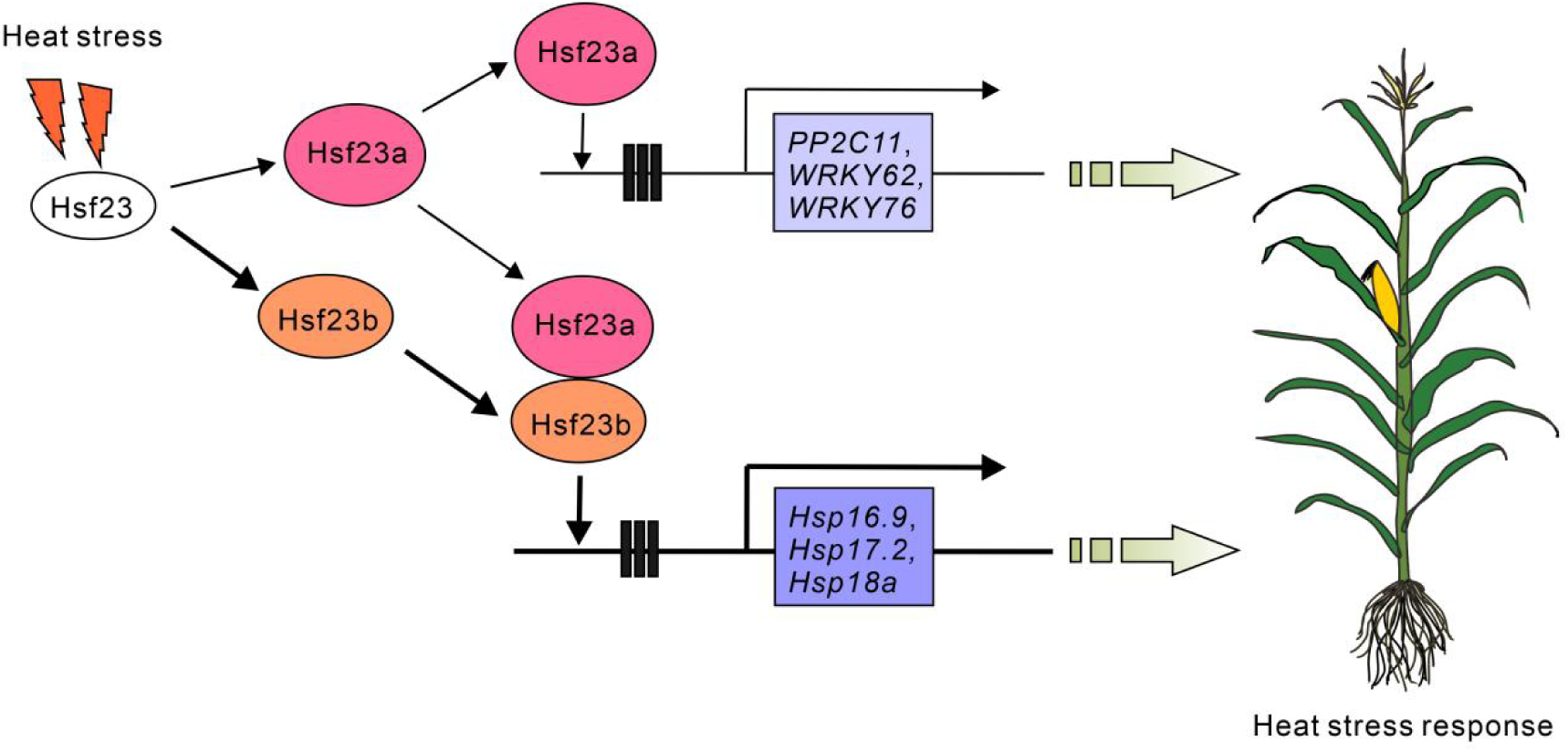
A proposed model for the role of Hsf23a and Hsf23b in regulating maize heat stress tolerance. Under HS conditions, the HS-induced AS of *Hsf23* produces *Hsf23a* and *Hsf23b* transcripts. Hsf23a directly activates the expression of *PP2C11* and *WRKYs* under heat stress. Similarly, Hsf23b directly activates *sHSP* expression. Additionally, Hsf23a interacts with Hsf23b to enhance the expression of Hsf23b-regulated *sHSP* genes, and strengthen the heat stress responses. The arrows represent positive effects and the thickness of the lines represents the degree of effects.

## Acknowledgements

This work was supported by the National Key Research and Development Program of China (2021YFF1000301) and the National Natural Science Foundation of China (31771805).

## Author contributions

JW, H-YJ and B-JC designed the experiments. JW analyzed the data and wrote the manuscript. JW, N-NS, Q-QQ and A-QS performed the experiments. W-NS performed the bioinformatics analysis. JW and H-YJ revised the manuscript. All authors read and approved of the manuscript.

## Data availability

The RNA-seq datasets from this article can be found in the National Center for Biotechnology Information (http://www.ncbi.nlm.nih.gov/) under accession number PRJNA1101208. The data that support the findings of this study are available in the Supporting Information.

## Supporting Information

Additional Supporting Information may be found online in the Supporting Information section at the end of the article.

**Fig. S1** Protein sequence alignment and phylogenetic relationships of Hsf23a and Hsf23b.

**Fig. S2** Phenotypes under normal conditions and DAB staining of the *Hsf23a*-overexpressing and *Hsf23b*-overexpressing plants.

**Fig. S3** Thermotolerance analysis of the transgenic Arabidopsis plants.

**Fig. S4** Identification of *hsf23a* and *hsf23a23b* mutants.

**Fig. S5** Differentially expressed genes (DEGs) in response to HS for WT, *OE23a* and *OE23b* plants.

**Fig. S6** GO terms for DEGs in the comparison between *OE23a, OE23b* and WT under normal growth conditions.

**Fig. S7** Transactivation assay of Hsf23a and Hsf23b.

**Table S1** Primers used in this study.

**Table S2** Differentially expressed genes (DEGs) in WT, *OE23a* and *OE23b* plants with and without heat treatment.

**Table S3** Differentially expressed genes (DEGs) in *OE23a* or *OE23b* compared with the WT plants at normal and heat stress conditions.

**Table S4** Hsf23a-regulated and Hsf23b-regulated *HR* genes.

**Table S5** The 11 Hsf23a-dependent and 371 Hsf23b-dependent *HR* genes.

## References

Andrási N, Pettkó-Szandtner A, Szabados L. 2021. Diversity of plant heat shock factors: regulation, interactions, and functions. Journal of Experimental Botany 72: 1558–1575.

Ashraf MF, Yang S, Wu R, Wang Y, Hussain A, Noman A, Khan MI, Liu Z, Qiu A, Guan D, et al. 2018. Capsicum annuum HsfB2a Positively Regulates the Response to Ralstonia solanacearum Infection or High Temperature and High Humidity Forming Transcriptional Cascade with CaWRKY6 and CaWRKY40. Plant and Cell Physiology 59: 2608–2623.

Begcy K, Nosenko T, Zhou LZ, Fragner L, Weckwerth W, Dresselhaus T. 2019. Male sterility in maize after transient heat stress during the tetrad stage of pollen development. Plant Physiology 181: 683–700.

Cao JM, Yao DM, Lin F, Jiang MY. 2014. PEG-mediated transient gene expression and silencing system in maize mesophyll protoplasts: a valuable tool for signal transduction study in maize. Acta Physiologiae Plantarum 36: 1271–1281.

Chan-Schaminet KY, Baniwal SK, Bublak D, Nover L, Scharf KD. 2009. Specific interaction between tomato HsfA1 and HsfA2 creates hetero-oligomeric superactivator complexes for synergistic activation of heat stress gene expression. Journal of Biological Chemistry 284: 20848–20857.

Chen H, Lai Z, Shi J, Xiao Y, Chen Z, Xu X. 2010. Roles of arabidopsis WRKY18, WRKY40 and WRKY60 transcription factors in plant responses to abscisic acid and abiotic stress. BMC Plant Biology 10: 281.

Cheng Q, Zhou Y, Liu Z, Zhang L, Song G, Guo Z, Wang W, Qu X, Zhu Y, Yang D. 2015. An alternatively spliced heat shock transcription factor, OsHSFA2dI, functions in the heat stress-induced unfolded protein response in rice. Plant Biology 17: 419–429.

Cicero MP, Hubl ST, Harrison CJ, Littlefield O, Hardy JA, Nelson HC. 2001. The wing in yeast heat shock transcription factor (HSF) DNA-binding domain is required for full activity. Nucleic Acids Research 29: 1715–1723.

Feng XH, Zhang HX, Ali M, Gai WX, Cheng GX, Yu QH, Yang SB, Li XX, Gong ZH. 2019. A small heat shock protein CaHsp25.9 positively regulates heat, salt, and drought stress tolerance in pepper (Capsicum annuum L.). Plant Physiology and Biochemistry 142: 151–162.

Fragkostefanakis S, Röth S, Schleiff E, Scharf KD. 2015. Prospects of engineering thermotolerance in crops through modulation of heat stress transcription factor and heat shock protein networks. Plant Cell and Environment 38: 1881–1895.

Fragkostefanakis S, Simm S, El-Shershaby A, Hu Y, Bublak D, Mesihovic A, Darm K, Mishra SK, Tschiersch B, Theres K et al. 2019. The repressor and co-activator HsfB1 regulates the major heat stress transcription factors in tomato. Plant Cell and Environment 42: 874–890.

Friedrich T, Oberkofler V, Trindade I, Altmann S, Brzezinka K, Lämke J, Gorka M, Kappel C, Sokolowska E, Skirycz A et al. 2019. Heteromeric HSFA2/HSFA3 complexes drive transcriptional memory after heat stress in Arabidopsis. Nature Communications 12: 3426.

Gu L, Jiang T, Zhang C, Li X, Wang C, Zhang Y, Li T, Dirk LMA, Downie AB, Zhao T. 2019. Maize HSFA2 and HSBP2 antagonistically modulate raffinose biosynthesis and heat tolerance in Arabidopsis. Plant Journal 100: 128–142.

Guo M, Liu JH, Ma X, Luo DX, Gong ZH, Lu MH. 2016. The plant heat stress transcription factors (HSFs): structure, regulation, and function in response to abiotic stresses. Frontiers in Plant Science 7: 114.

Hu W, Ren Q, Chen Y, Xu G, Qian Y. 2021. Genome-wide identification and analysis of WRKY gene family in maize provide insights into regulatory network in response to abiotic stresses. BMC Plant Biology 21: 427.

Hu Y, Mesihovic A, Jiménez-Gómez JM, Röth S, Gebhardt P, Bublak D, Bovy A, Scharf KD, Schleiff E, Fragkostefanakis S. 2020. Natural variation in HsfA2 pre-mRNA splicing is associated with changes in thermotolerance during tomato domestication. New Phytologist 225: 1297–1310.

Huang YC, Niu CY, Yang CR, Jinn TL. 2016. The Heat Stress Factor HSFA6b Connects ABA Signaling and ABA-Mediated Heat Responses. Plant Physiology 172: 1182–1199.

Hwang SM, Kim DW, Woo MS, Jeong HS, Son YS, Akhter S, Choi GJ, Bahk JD. 2014. Functional characterization of Arabidopsis HsfA6a as a heat-shock transcription factor under high salinity and dehydration conditions. Plant Cell and Environment 37: 1202–1222.

Jiang J, Liu X, Liu C, Liu G, Li S, Wang L. 2017. Integrating Omics and Alternative Splicing Reveals Insights into Grape Response to High Temperature. Plant Physiology 173: 1502–1518.

Komatsu K, Suzuki N, Kuwamura M, Nishikawa Y, Nakatani M, Ohtawa H, Takezawa D, Seki M, Tanaka M, Taji T et al. 2013. Group A PP2Cs evolved in land plants as key regulators of intrinsic desiccation tolerance. Nature Communications 4: 2219.

Laloum T, Martín G, Duque P. 2018. Alternative Splicing Control of Abiotic Stress Responses. Trends in Plant Science 23: 140–150.

Lesk C, Rowhani P, Ramankutty N. 2016. Influence of extreme weather disasters on global crop production. Nature 529: 84–87.

Li B, Gao K, Ren H, Tang W. 2018. Molecular mechanisms governing plant responses to high temperatures. Journal of Integrative Plant Biology 60: 757–779.

Li Z, Tang J, Bassham DC, Howell SH. 2021. Daily temperature cycles promote alternative splicing of RNAs encoding SR45a, a splicing regulator in maize. Plant Physiology 186: 1318–1335.

Li Z, Tang J, Srivastava R, Bassham DC, Howell SH. 2020. The Transcription Factor bZIP60 Links the Unfolded Protein Response to the Heat Stress Response in Maize. Plant Cell 32: 3559–3575.

Lin J, Xu Y, Zhu Z. 2020. Emerging plant thermosensors: from RNA to protein. Trends in Plant Science 25: 1187–1189.

Ling Y, Mahfouz MM, Zhou S. 2021. Pre-mRNA alternative splicing as a modulator for heat stress response in plants. Trends in Plant Science 26: 1153–1170.

Liu HC, Charng YY. 2013. Common and distinct functions of Arabidopsis class A1 and A2 heat shock factors in diverse abiotic stress responses and development. Plant Physiology 163: 276–290.

Liu HC, Liao HT, Charng YY. 2011. The role of class A1 heat shock factors (HSFA1s) in response to heat and other stresses in Arabidopsis. Plant Cell and Environment 34: 738–751.

Liu J, Sun N, Liu M, Liu J, Du B, Wang X, Qi X. 2013. An autoregulatory loop controlling Arabidopsis HsfA2 expression: role of heatshock-induced alternative splicing. Plant Physiology 162: 512–521.

Lohani N, Singh MB, Bhalla PL. 2020. High temperature susceptibility of sexual reproduction in crop plants. Journal of Experimental Botany 71: 555–568.

Long Y, Qin Q, Zhang J, Zhu Z, Liu Y, Gu L, Jiang H, Si W. 2023. Transcriptomic and weighted gene co-expression network analysis of tropic and temperate maize inbred lines recovering from heat stress. Plant Science 327: 111538.

McLoughlin F, Basha E, Fowler ME, Kim M, Bordowitz J, Katiyar-Agarwal S, Vierling E. 2016. Class I and II small heat shock proteins together with HSP101 protect protein translation factors during heat stress. Plant Physiology 172: 1221–1236.

Miao J, Li X, Li X, Tan W, You A, Wu S, Tao Y, Chen C, Wang J, Zhang D et al. 2020. OsPP2C09, a negative regulatory factor in abscisic acid signalling, plays an essential role in balancing plant growth and drought tolerance in rice. New Phytologist 227: 1417–1433.

Moore CE, Meacham-Hensold K, Lemonnier P, Slattery RA, Benjamin C, Bernacchi CJ, Lawson T, Cavanagh AP. 2021. The effect of increasing temperature on crop photosynthesis: from enzymes to ecosystems. Journal of Experimental Botany 72: 2822–2844.

Mou S, He W, Jiang H, Meng Q, Zhang T, Liu Z, Qiu A, He S. 2024. Transcription factor CaHDZ15 promotes pepper basal thermotolerance by activating HEAT SHOCK FACTORA6a. Plant Physiology 25: kiae037.

Ohama N, Sato H, Shinozaki K, Yamaguchi-Shinozaki K. 2017. Transcriptional Regulatory Network of Plant Heat Stress Response. Trends in Plant Science 22: 53–65.

Samtani H, Sharma A, Khurana P. 2023. Ectopic overexpression of TaHsfA5 promotes thermomorphogenesis in Arabidopsis thaliana and thermotolerance in Oryza sativa. Plant Molecular Biology 112: 225–243.

Slafer GA, Savin R. 2018. Can N management affect the magnitude of yield loss due to heat waves in wheat and maize? Current Opinion in Plant Biology 45: 276–283.

Wang YX, Yu TF, Wang CX, Wei JT, Zhang SX, Liu YW, Chen J, Zhou YB, Chen M, Ma YZ et al. 2023. Heat shock protein TaHSP17.4, a TaHOP interactor in wheat, improves plant stress tolerance. International Journal of Biological Macromolecules 246: 125694.

Wei K, Pan S. 2014. Maize protein phosphatase gene family: identification and molecular characterization. BMC Genomics 15: 773.

Wen J, Qin Z, Sun L, Zhang Y, Wang D, Peng H, Yao Y, Hu Z, Ni Z, Sun Q et al. 2023. Alternative splicing of TaHSFA6e modulates heat shock protein-mediated translational regulation in response to heat stress in wheat. New Phytologist 239: 2235–2247.

Wu Z, Li T, Ding L, Wang C, Teng R, Xu S, Cao X, Teng N. 2024. Lily LlHSFC2 coordinates with HSFAs to balance heat stress response and improve thermotolerance. New Phytologist 241: 2124–2142.

Wu Z, Liang J, Wang C, Ding L, Zhao X, Cao X, Xu S, Teng N, Yi M. 2019. Alternative Splicing Provides a Mechanism to Regulate LlHSFA3 Function in Response to Heat Stress in Lily. Plant Physiology 181: 1651–1667.

Wu Z, Liang J, Wang C, Zhao X, Zhong X, Cao X, Li G, He J, Yi M. 2018. Overexpression of lily HsfA3s in Arabidopsis confers increased thermotolerance and salt sensitivity via alterations in proline catabolism. Journal of Experimental Botany 69: 2005–2021.

Xin H, Zhang H, Chen L, Li X, Lian Q, Yuan X, Hu X, Cao L, He X, Yi M. 2010. Cloning and characterization of HsfA2 from Lily (Lilium longiflorum). Plant Cell Reports 29: 875–885.

Xue GP, Drenth J, McIntyre CL. 2015. TaHsfA6f is a transcriptionalactivator that regulates a suite of heat stress protection genes in wheat (Triticumaestivum L.) including previously unknown Hsf targets. Journal of Experimental Botany 66: 1025–1039.

Zhang H, Li G, Fu C, Duan S, Hu D, Guo X. 2020. Genome-wide identification, transcriptome analysis and alternative splicingevents of Hsf family genes in maize. Scientific Reports 10: 8073.

Zhao C, Liu B, Piao S, Wang X, Lobell DB, Huang Y, Huang M, Yao Y, Bassu S, Ciais P et al. 2017. Temperature increase reduces global yields of major crops in four independent estimates. *Proceedings of the National Academy of Sciences*, USA 114: 9326–9331.

