## Supplemental figures for "The alternative splicing of *ZmHsf23* regulates heat stress tolerance in maize"

***New Phytologist* Supporting Information**

The following Supporting Information is available for this article:

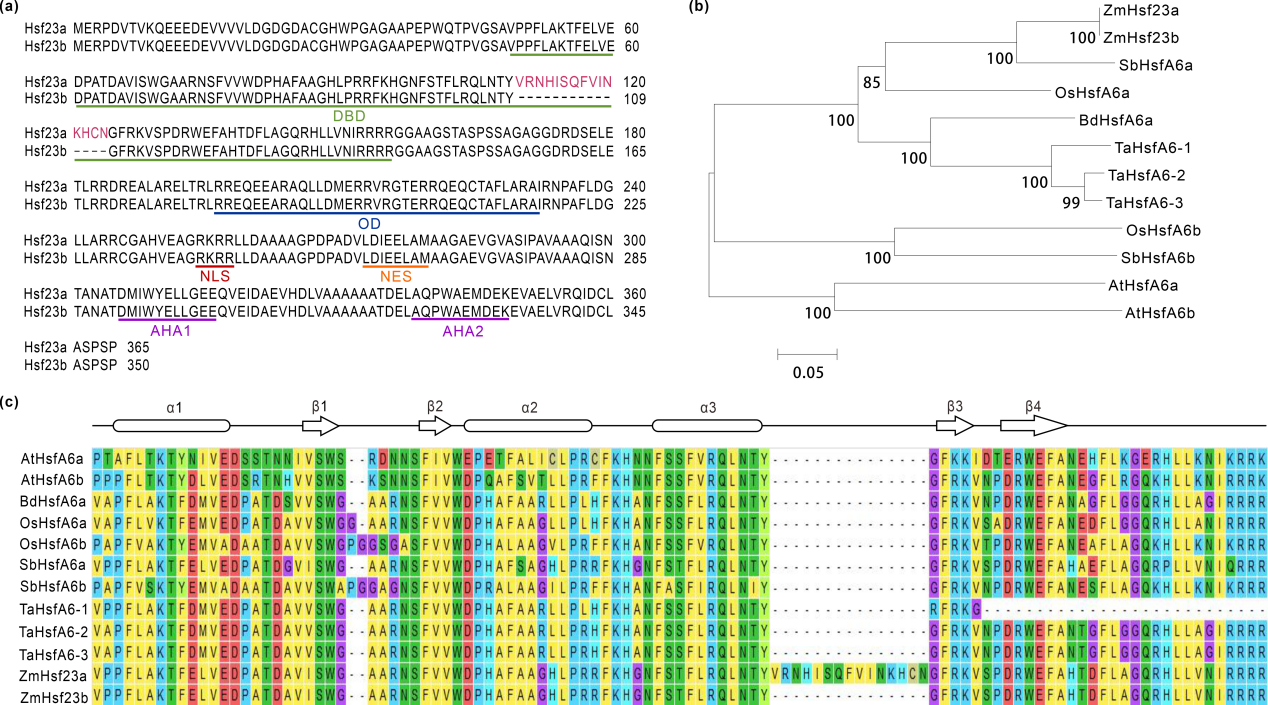

**Fig. S1** Protein sequence alignment and phylogenetic relationships of Hsf23a and Hsf23b. (a), Protein sequence alignment of Hsf23a and Hsf23b. The solid lines with different colors indicate different domains. The red amino acids are encoded by the miniexon. (b), Phylogenetic analysis of HsfA6 proteins from different plants. Zm, *Zea mays*; At, *Arabidopsis thaliana*; Os, *Oryza sativa*; Bd, *Brachypodium distachyon*; Ta, *Triticum aestivum*; Sb, *Sorghum bicolor*. The phylogenetic tree was constructed in MEGA 7 using the neighbor-joining method. (c), Multiple sequence alignment of the DBD domain of HsfA6s in different plants. The secondary structure elements of DBD (α1-β1-β2-α2-α3-β3-β4) are shown with arrows and rounded rectangle above the alignment.

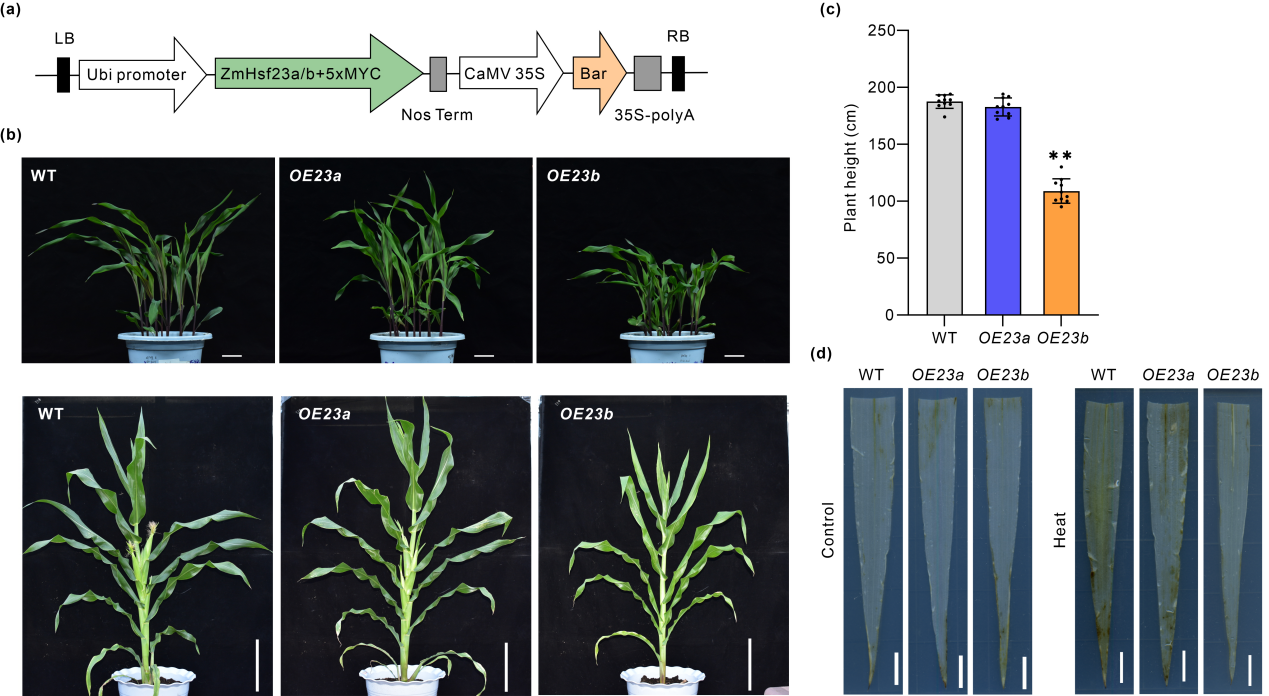

**Fig. S2** Phenotypes under normal conditions and DAB staining of the *Hsf23a*-overexpressing and *Hsf23b*-overexpressing plants. (a), Schematic diagram of *Hsf23a* and *Hsf23b* overexpression vector construction. (b), Phenotypes of WT and transgenic seedlings (first row) and mature plants (second row) under normal growth conditions. Scale bars, 5 cm (first row) and 20 cm (second row). (c), Comparison of plant height between WT, *OE23a* and *OE23b* plants at the mature stage. Bars represent means ± SD (n=10 repeats). Significant differences (Student’s t test: **P<0.01). (d), H_2_O_2_ accumulation was detected with DAB in maize leaves before and after heat treatment. Leaves from WT, *OE23a* and *OE23b* seedlings before and after 45°C HS for 10 h, were stained. Scale bar=1 cm.

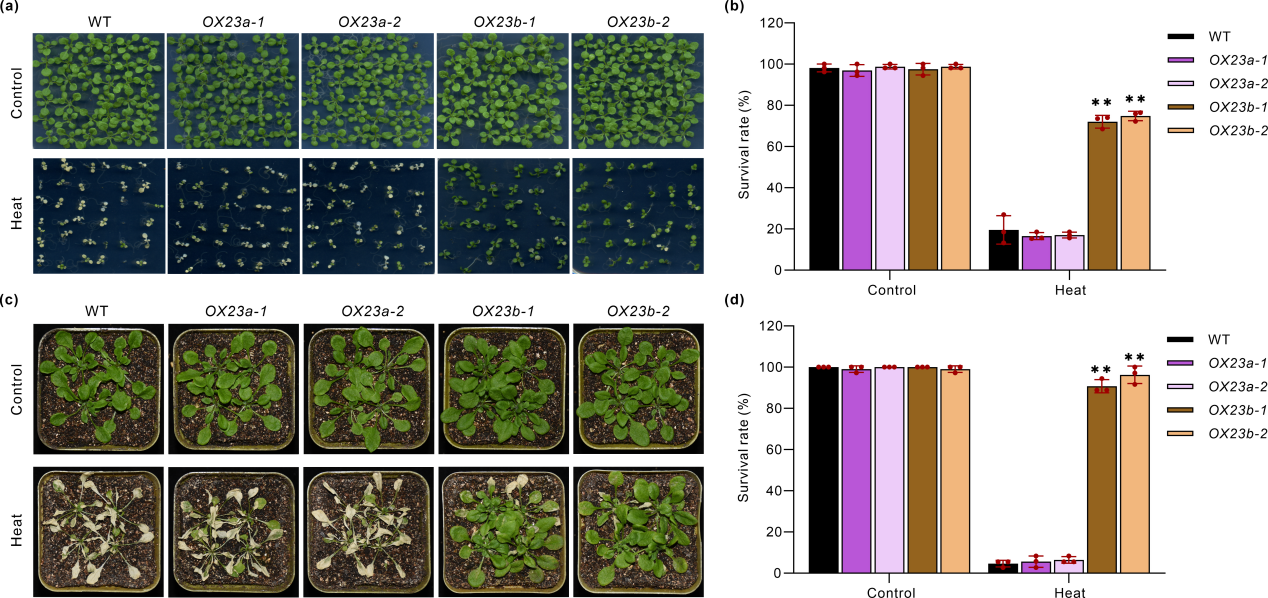

**Fig. S3** Thermotolerance analysis of the transgenic Arabidopsis plants. (a), Heat phenotypes of WT, *ZmHsf23a* transgenic (*OX23a-1*,*2*) and *ZmHsf23b* transgenic (*OX23b-1*,*2*) seedlings during seed germination. The WT and transgenic plants were germinated on MS media for 5 d at normal conditions and then treated with or without 45°C for 1.5 h. Pictures were taken after recovery for 3 days. (b), The survival rate of seedlings in (a). Bars represent means ± SD (n = 3 repeats). (Student’s t test: **P < 0.01). (c, d), Heat phenotypes (c) and survival rates (d) of WT and transgenic seedlings before and after heat treatment. Two-week-old seedlings of WT, *ZmHsf23a* transgenic and *ZmHsf23b* transgenic were treated with or without 45°C for 5.5 h. Pictures were taken after recovery for 3 days. Bars represent means ± SD (n = 3 repeats). (Student’s t test: **P < 0.01).

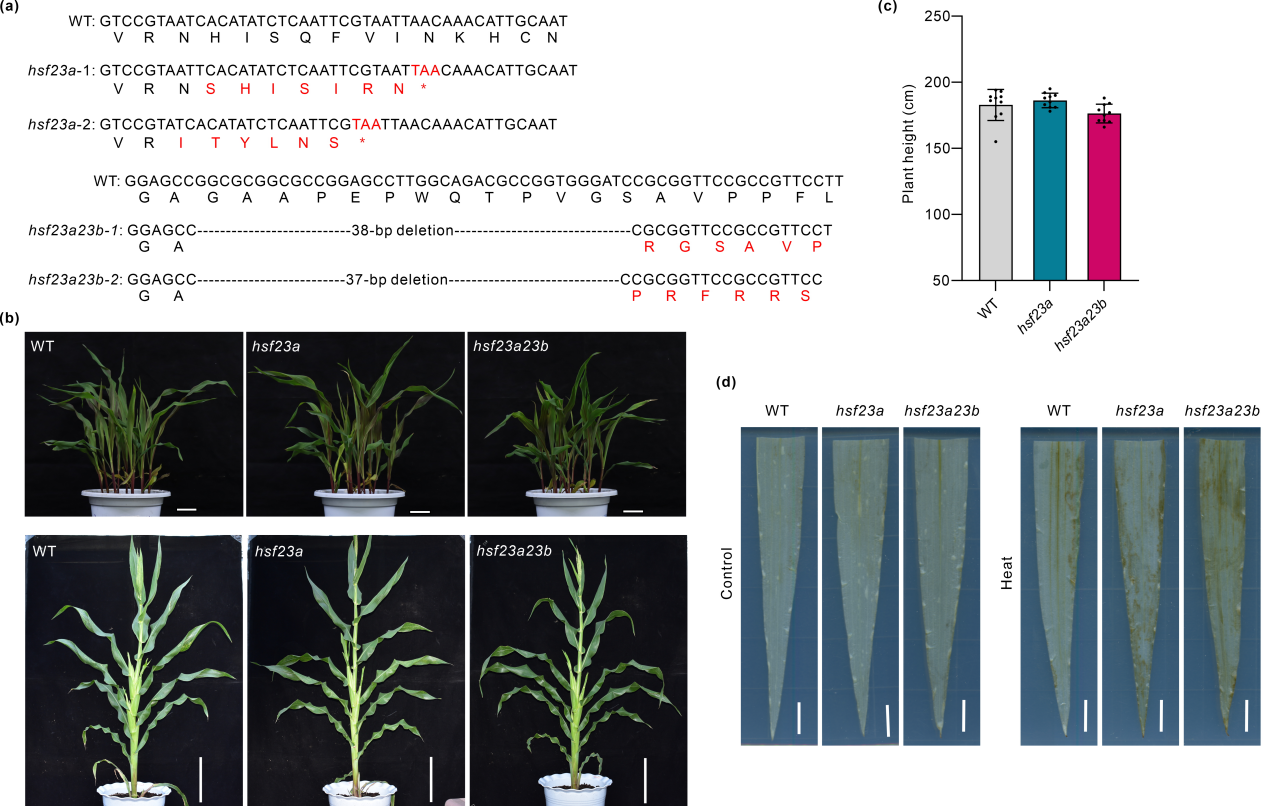

**Fig. S4** Identification of *hsf23a* and *hsf23a23b* mutants. (a), The nucleic acid and deduced amino acid sequences of WT, *hsf23a* and *hsf23a23b*. The red font indicates the mutated amino acid sequences. (b), Phenotype of WT, *hsf23a* and *hsf23a23b* mutants at the seedlings stage (first row) and mature stage (second row) under normal growth conditions. Scale bars, 5 cm (first row) and 20 cm (second row). (c), Comparison of plant height between WT, *hsf23a* and *hsf23a23b* plants at the mature stage. Bars represent means ± SD (n=10 repeats). Significant differences (Student’s t test: **P<0.01). (d), H_2_O_2_ accumulation was detected with DAB in maize leaves before and after heat treatment. Leaves from WT, *hsf23a* and *hsf23a23b* seedlings before and after 45°C HS for 6 h, were stained. Scale bar=1 cm.

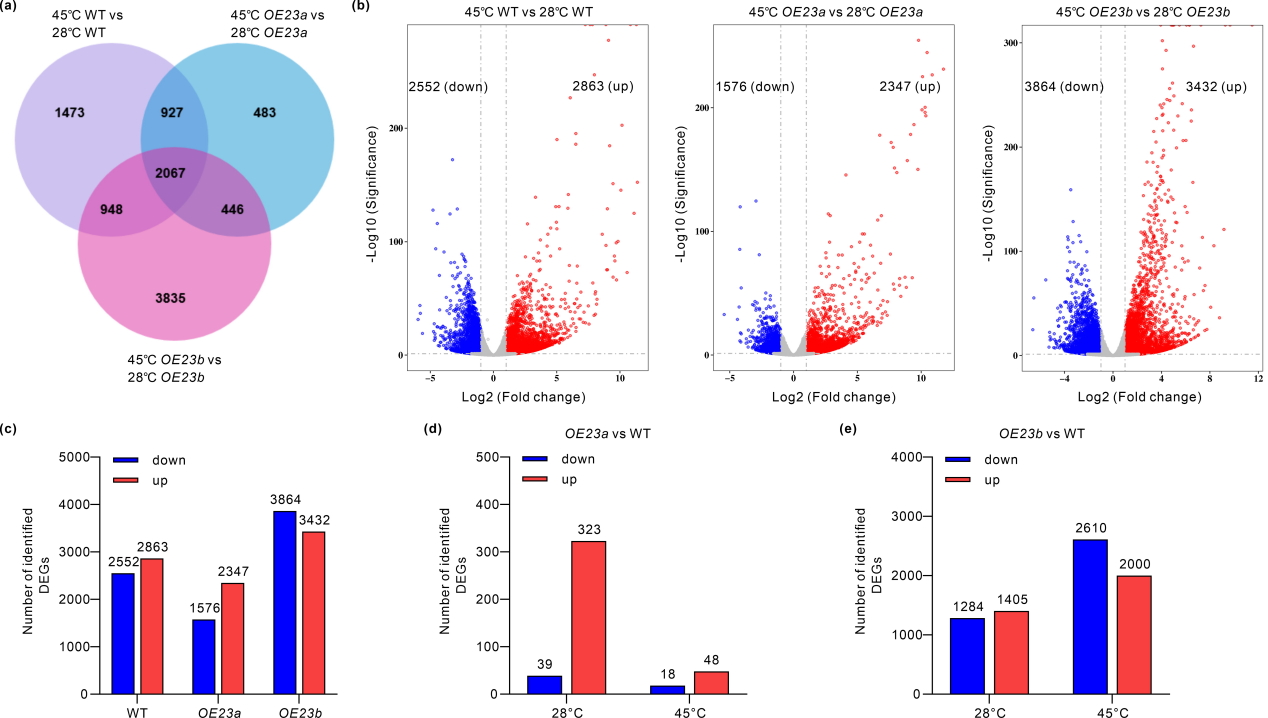

**Fig. S5** Differentially expressed genes (DEGs) in response to HS for WT, *OE23a* and *OE23b* plants. (a), Venn diagrams representing the overlap between DEGs in the WT, *OE23a* and *OE23b* plants in response to HS. (b), Volcano plots represent the number of DEGs that were upregulated or downregulated in response to HS. (c), Histogram showing genes upregulated or downregulated in WT, *OE23a* and *OE23b* plants in response to HS. Numbers on the top indicate the number of DEGs in the comparison. (d), DEGs in the comparison between *OE23a* and WT at 28°C and 45°C. (e), DEGs in the comparison between *OE23b* and WT in response to HS. DEGs were identified with p < 0.05 and absolute log2 fold value ≥ 1.

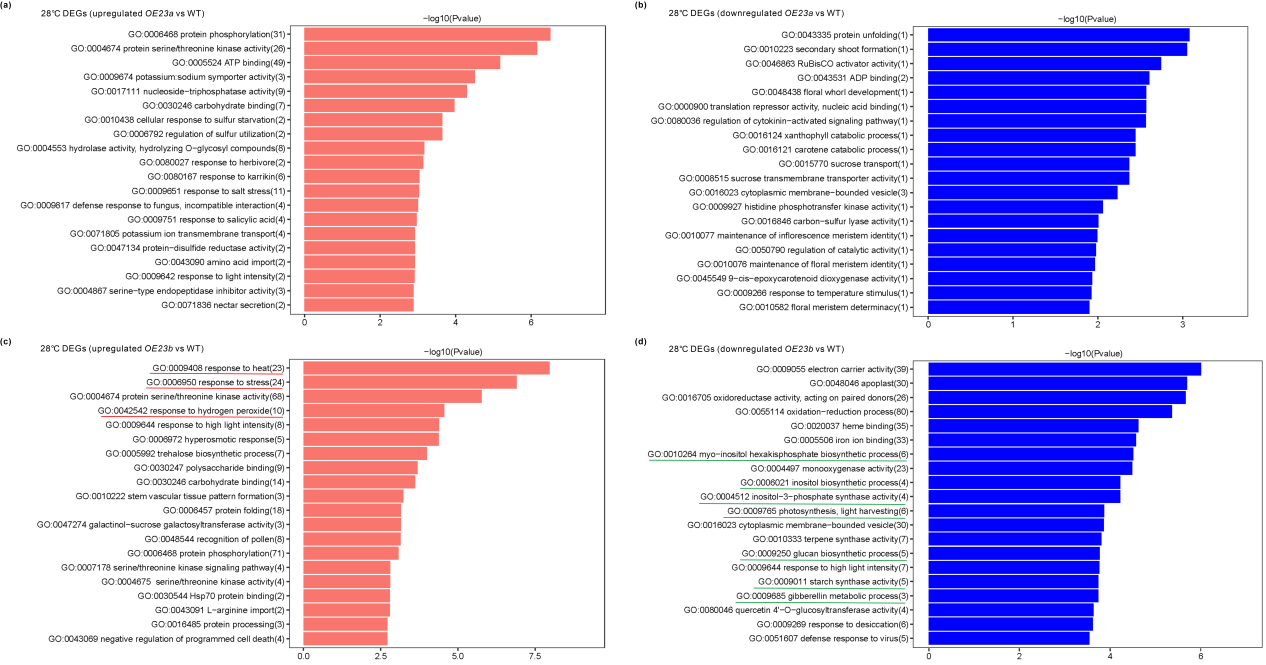

**Fig. S6** GO terms for DEGs in the comparison between *OE23a*, *OE23b* and WT under normal growth conditions. (a, b), GO terms for upregulated (a) and downregulated (b) genes in the comparison between *OE23a* and WT. (c, d), GO terms for upregulated (c) and downregulated (d) genes in the comparison between *OE23b* and WT.

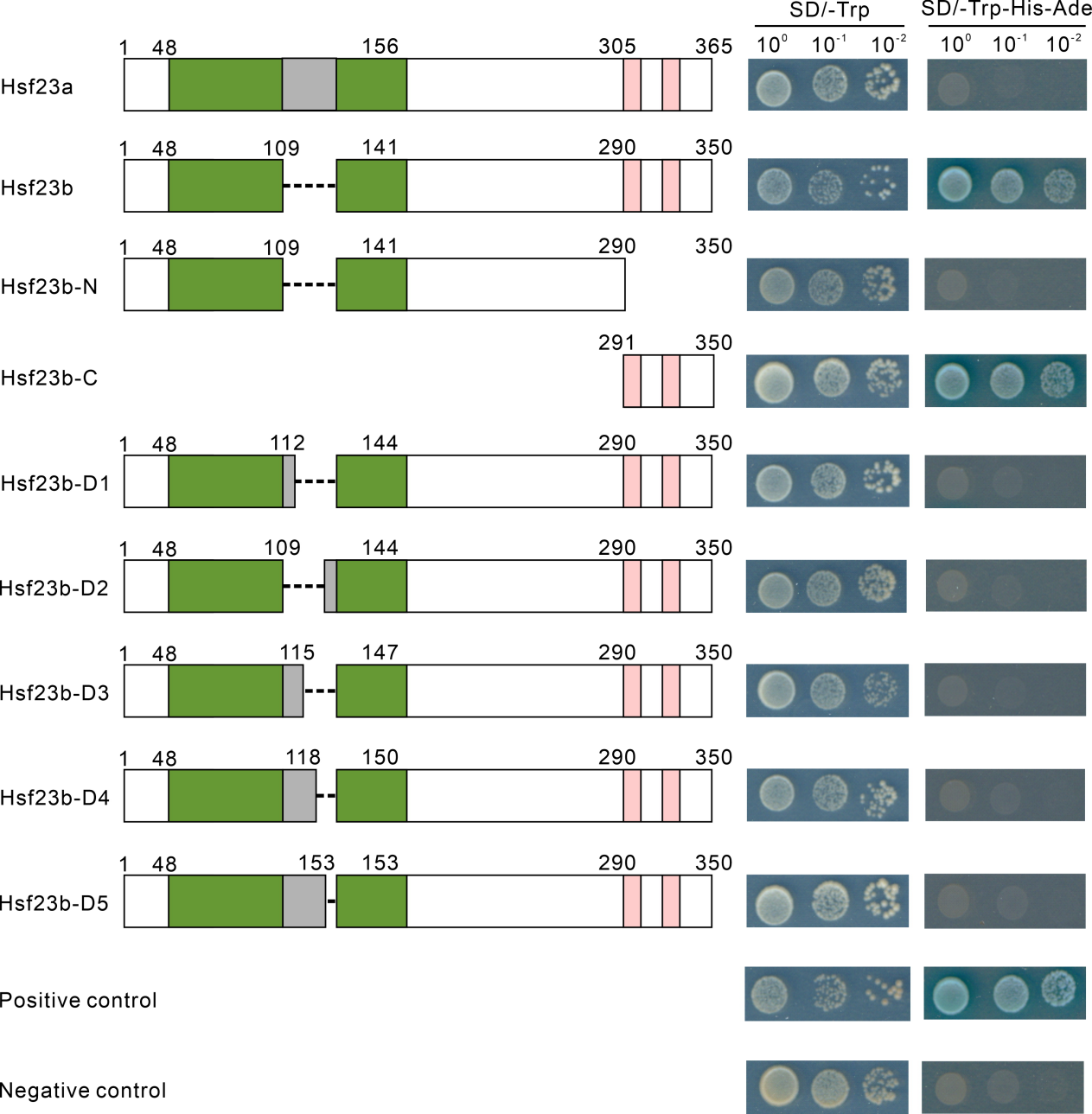

**Fig. S7** Transactivation assay of Hsf23a and Hsf23b. The structures of two splice variant proteins and multiple wing truncated Hsf23b constructions (left) were used for transactivation activity assay. Green box indicates the DBD domain. Gray box indicates the wing sequences. Different vectors were separately transformed into the yeast cells.
